# Ongoing Global and Regional Adaptive Evolution of SARS-CoV-2

**DOI:** 10.1101/2020.10.12.336644

**Authors:** Nash D. Rochman, Yuri I. Wolf, Guilhem Faure, Pascal Mutz, Feng Zhang, Eugene V. Koonin

## Abstract

Understanding the trends in SARS-CoV-2 evolution is paramount to control the COVID- 19 pandemic. We analyzed more than 300,000 high quality genome sequences of SARS-CoV-2 variants available as of January 2021. The results show that the ongoing evolution of SARS-CoV-2 during the pandemic is characterized primarily by purifying selection, but a small set of sites appear to evolve under positive selection. The receptor-binding domain of the spike protein and the nuclear localization signal (NLS) associated region of the nucleocapsid protein are enriched with positively selected amino acid replacements. These replacements form a strongly connected network of apparent epistatic interactions and are signatures of major partitions in the SARS-CoV-2 phylogeny. Virus diversity within each geographic region has been steadily growing for the entirety of the pandemic, but analysis of the phylogenetic distances between pairs of regions reveals four distinct periods based on global partitioning of the tree and the emergence of key mutations. The initial period of rapid diversification into region-specific phylogenies that ended in February 2020 was followed by a major extinction event and global homogenization concomitant with the spread of D614G in the spike protein, ending in March 2020. The NLS associated variants across multiple partitions rose to global prominence in March-July, during a period of stasis in terms of inter-regional diversity. Finally, beginning July 2020, multiple mutations, some of which have since been demonstrated to enable antibody evasion, began to emerge associated with ongoing regional diversification, which might be indicative of speciation.

**Significance:** Understanding the ongoing evolution of SARS-CoV-2 is essential to control and ultimately end the pandemic. We analyzed more than 300,000 SARS-CoV-2 genomes available as of January 2021 and demonstrate adaptive evolution of the virus that affects, primarily, multiple sites in the spike and nucleocapsid protein. Selection appears to act on combinations of mutations in these and other SARS-CoV-2 genes. Evolution of the virus is accompanied by ongoing adaptive diversification within and between geographic regions. This diversification could substantially prolong the pandemic and the vaccination campaign, in which variant-specific vaccines are likely to be required.

## Introduction

High mutation rates of RNA viruses enable adaptation to hosts at a staggering pace (1–4). Nevertheless, robust sequence conservation indicates that purifying selection is the principal force shaping the evolution of virus populations, with positive selection affecting only relatively small subsets of sites directly involved in virus-host coevolution (5–8). The fate of a novel zoonotic virus is in part determined by the race between public health intervention and virus diversification. Even intermittent periods of positive selection can result in lasting immune evasion, leading to oscillations in the size of the susceptible population, and ultimately, a regular pattern of repeating epidemics, as has been amply demonstrated for Influenza(9–11).

During the current coronavirus pandemic (COVID-19), understanding the degree and dynamics of the diversification of severe acute respiratory syndrome coronavirus 2 (SARS-Cov-2) and identification of sites subject to positive selection are essential for establishing practicable, proportionate public health responses, from guidelines on isolation and quarantine to vaccination(12). To investigate the evolution of SARS-CoV- 2, we collected all available SARS-Cov-2 genomes as of January 8, 2021, and constructed a global phylogenetic tree using a “divide and conquer” approach. Patterns of repeated mutations fixed along the tree were analyzed in order to identify the sites subject to positive selection. These sites form a network of potential epistatic interactions. Analysis of the putative adaptive mutations provides for the identification of signatures of evolutionary partitions of SARS-CoV-2. The dynamics of these partitions over the course of the pandemic reveals alternating periods of globalization and regional diversification.

## Results and Discussion

### Global multiple sequence alignment of the SARS-CoV-2 genomes

To investigate the evolution of SARS-CoV-2, we aggregated all available SARS-Cov-2 genomes as of January 8, 2021, from the three principal repositories: Genbank(13), Gisaid(14), and CNCB(15). From the total of 321,096 submissions in these databases, 175,857 unique SARS-Cov-2 genome sequences were identified, and 98,185 high quality sequences were incorporated into a global multisequence alignment (MSA) consisting of the concatenated open reading frames with stop codons trimmed. The vast majority of the sequences excluded from the MSA were removed due to a preponderance of ambiguous characters (see Methods). The sequences in the final MSA correspond to 175,776 isolates with associated date and location metadata.

### Tree Construction

Several methods for coronavirus phylogenetic tree inference have been tested(16, 17). The construction of a single high-quality tree from nearly 200,000 30 kilobase (kb) sequences using any of the existing advanced methods is computationally prohibitive. Iterative construction of the complete phylogeny would seem an obvious solution such that a global topology would be obtained based on a subset of sequences available at an earlier date, and later sequences would be incorporated into the existing tree.

However, this approach induces artifacts through the inheritance of deep topologies that differ substantially from any maximum-likelihood solution corresponding to the complete alignment.

Therefore, building on the available techniques, we utilized a “divide and conquer” approach which is not subject to these artifacts and furthermore can be employed for datasets that cannot be structured by sequencing date, including metagenomic analyses. This approach leverages two ideas. First, for any alignment, a diverse representative subset of sequences can be used to establish a deep topology, the tree “skeleton”, that corresponds to a maximum-likelihood solution over the entire alignment. Second, deep branches in an unrooted tree are primarily determined by common substitutions relative to consensus. In other words, rare substitutions are unlikely to affect deep splits or branch lengths. We adopted the following steps to resolve the global phylogeny for SARS-CoV-2 (see Methods for details).

The first step is to construct sets of diverse representative sequences such that the topologies inferred from each subset share the same tree “skeleton” corresponding to that of the global topology. Sequence diversity is measured by the hamming distance between pairs of sequences; however, maximizing hamming distances among a set of representative sequences does not guarantee maximization of the tree distances in the resultant global topology and so does not guarantee the maximum-likelihood topology of this subset would share the global tree skeleton. Therefore, a reduced alignment containing only the top 5% of sites (ignoring nearly-invariant sites, see below) with the most common substitutions relative to consensus was constructed. Sequences redundant over this narrow alignment were removed. Sets of diverse, based on the hamming distances over this reduced alignment, representative sequences were then achieved and all subtrees generated from each diverse subset were aggregated to constrain a single, composite tree.

This composite tree reflects the correct tree skeleton and could be used to constrain the global topology; however, due to numerous sequencing errors in this dataset, another intermediate step was taken. A second reduced alignment was constructed in which nearly-invariant sites, which may represent sequencing errors and should not be used to infer tree topology, were omitted. As before, sequences redundant over this reduced alignment were removed and a tree was then constructed from this alignment, constrained to maintain the topology of the composite tree wherever possible. This tree reflects the correct topology of the global tree but has incorrect branch lengths. Finally, the global tree with the correct branch lengths (Fig. 1A) was constructed over the whole alignment, constrained to maintain the topology of previous tree wherever possible. A complete reconstruction of ancestral sequences was then performed by leveraging Fitch traceback(18), enabling comprehensive identification of nucleotide and amino acid replacements across the tree.

**Figure 1.**
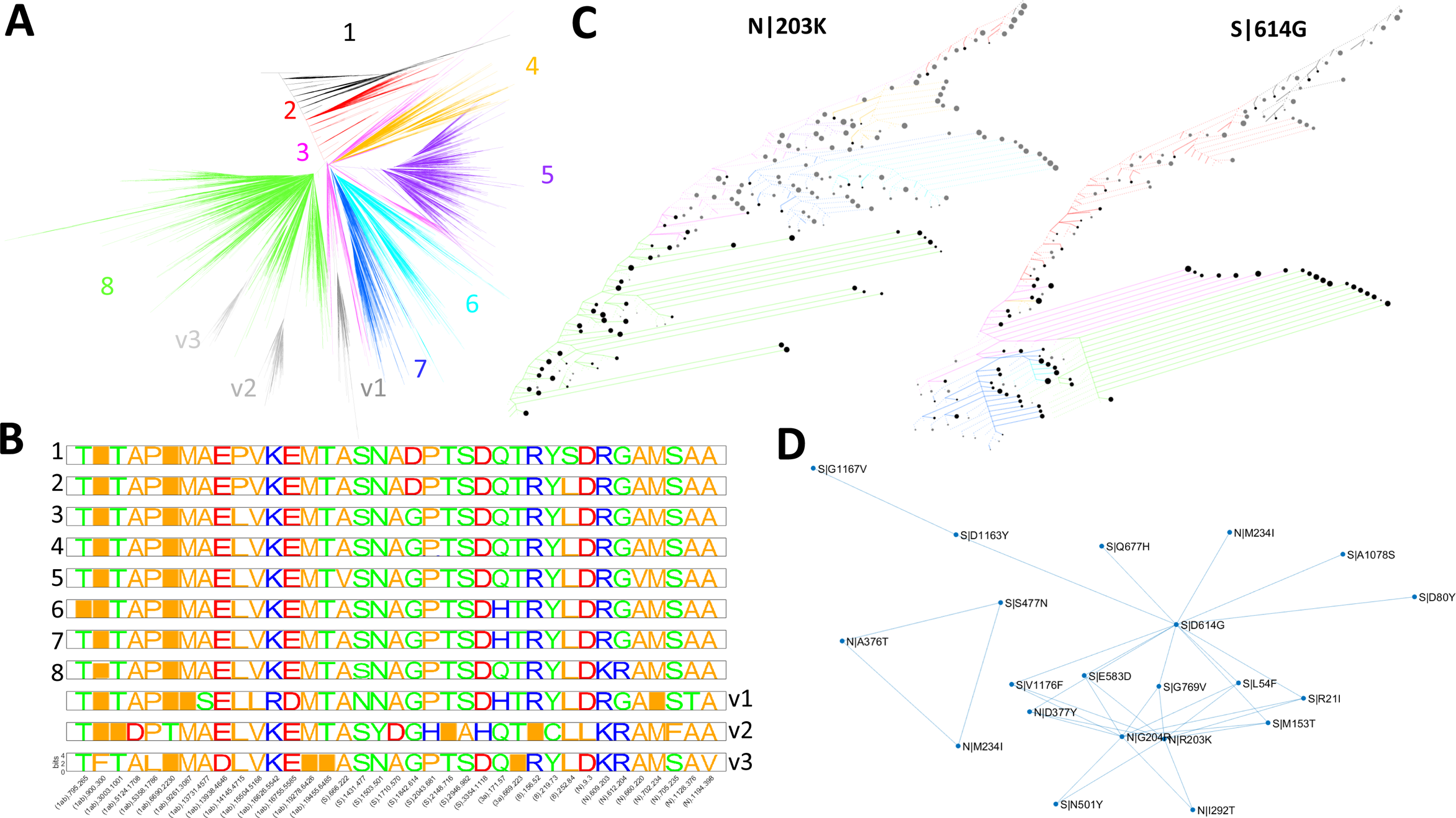
Evolution of SARS-CoV-2. **A**. Global tree reconstruction with 8 principal partitions and 3 variant clades enumerated and color-coded. **B**. Signatures of amino acid replacements for each partition. Sites are ordered as they appear in the genome. The proteins along with nucleotide and amino acid numbers are indicated underneath each column. **C**. Site history trees for spike 614 and nucleocapsid 203 positions. Nodes were included in this reduced tree based on the following criteria: those immediately succeeding a substitution; representing the last common ancestor of at least two substitutions; or terminal nodes representing branches of five sequences or more (approximately, based on tree weight). Edges are colored according to their position in the main partitions and the line type corresponds to the target mutation (solid) or any other state (dashed). Synonymous mutations are not shown. These sites are largely binary as are most sites in the genome. The sizes of the terminal node sizes are proportional to the log of the weight descendent from that node beyond which no substitutions in the site occurred. Node color corresponds to target mutation (black) or any other state (gray). **D**. Network of putative epistatic interactions for likely positively selected residues in the N and S proteins.

We identified 8 principal partitions within this tree, in a general agreement with other work(19–21), along with three divergent clades (Fig. 1A) that, as discussed below, are important for the interpretation of the metadata. Given the short evolutionary distances between SARS-CoV-2 isolates, despite the efforts described above, the topology of the global tree is a cause of legitimate concern(17, 22–24). For the analyses presented below, we rely on a single, explicit tree topology which is probably one of many equally likely estimates(17). Therefore, we sought to validate the robustness of the major partitions of the virus genomes using a phylogeny-free approach. To this end, pairwise Hamming distances were computed for all sequences in the MSA and the resulting distance matrix was embedded in a 3-dimensional subspace using classical multidimensional scaling. In this embedding, the 8 partitions are separated, and the optimal clustering, determined by *k*-means, returned 5 categories (see Methods, Fig. S1), of which 4 correspond to partitions 5 and 8, and the divergent clades v1 and v2. These findings indicate that an alternative tree with a comparable likelihood but a dramatically different coarse-grain topology, most likely, cannot be constructed from this MSA.

### Mutational Signatures&Biases and Estimation of Selection

Each of the 8 partitions and 3 variant clades can be characterized by a specific amino acid replacement signature (Fig. 1B), generally, corresponding to the most prominent amino acid replacements across the tree (Table S1), some of which are shared by two or more partitions and appear independently many times, consistent with other reports(25). The receptor binding domain (RBD) of the spike protein and a region of the nucleocapsid protein associated with nuclear localization signals (NLS)(26) are enriched with these signature replacements, but they are also found in the nonstructural proteins 1ab, 3a, and 8. The identification of these prevailing non-synonymous substitutions and an additional set of frequent synonymous substitutions suggested that certain sites in the SARS-CoV-2 genome might be evolving under positive selection. However, uncovering the selective pressures affecting virus evolution was complicated by non-negligible mutational biases. The distributions of the numbers of both synonymous and non-synonymous substitutions across the genome were found to be substantially over- dispersed compared to both the Poisson and normal expectations (Fig. S2).

Examination of the relative frequencies of all 12 possible nucleotide substitutions indicated a significant genome-wide excess of C to U mutations, approximately threefold higher than any other nucleotide substitution, with the exception of G to U, as well as some region-specific trends. Specifically, G to U mutations increase steadily in frequency throughout the second half of the genome and the distribution of nucleotide substitutions over the polyprotein is dramatically different from other ORFs (Fig. S3).

Motivated by the observation of the mutational biases, we compared the trinucleotide contexts of synonymous and non-synonymous substitutions as well as the contexts of low and high frequency substitutions. The contexts of high-frequency events, both synonymous and non-synonymous, were found to be dramatically different from the background frequencies. The NCN context (that is, all C->D mutations) harbors substantially more events than other contexts (all 16 NCN triplets are within the top 20 most high-frequency-biased, see Methods and Fig. S4) and is enriched in mutations uniformly across the genome, primarily, among high-frequency sites. This pattern suggests a mechanistic bias of the errors made by the coronavirus RNA-dependent RNA polymerase (RdRP). Evidently, such a bias that increases the likelihood of observing multiple, independent substitutions in the NCN context complicates the detection of selection pressures. However, only 2 of the 9 contexts with an excess of non-synonymous events are NCN (gct,tct, Fig. S4), suggesting that at least some of these repeated, non-synonymous mutations are driven by other mechanisms. Thus, we excluded all synonymous substitutions and non-synonymous substitutions with the NCN context from further consideration in the determination of candidate sites evolving under positive selection.

Beyond this specific context, the presence of any hypervariable sites complicates the computation of the *dN/dS* ratio, the gauge of protein-level selection(27), which requires enumerating the number of synonymous and non-synonymous substitutions within each gene. Hypervariable sites bias this analysis, and therefore, we used two methods to ensure reliable estimation of *dN/dS*. For each protein-coding gene of SARS-CoV-2 (splitting the long orf1ab into 15 constituent non-structural proteins), we obtained both a maximum likelihood estimate of *dN/dS* across 10 sub-alignments and an approximation computed from the global ancestral reconstruction (see Methods). This approach was required due to the size of the alignment, which makes a global maximum likelihood estimation computationally prohibitive. Despite considerable variability among the genes, we obtained estimates of substantial purifying selection (0.1<*dN/dS*<0.5) across most of the genome(Fig. S5), with a reasonable agreement between the two methods. This estimate is compatible with previous demonstrations of purifying selection affecting about 50% of the sites surveyed or more(5), among diverse RNA viruses(6).

### Evidence of Positive Selection

As shown in the previous section, evolution of SARS-CoV-2 is likely primarily driven by substantial purifying selection. However, more than 100 non-synonymous substitutions appeared to have emerged multiple times, independently, covering a substantial portion of the tree equivalent to approximately 200 or more terminal branches or “leaves”, and were not subject to an overt mechanistic bias. Due to the existence of many equally likely trees, in principle, in one or more of such trees, any of these mutations could resolve to a single event. However, such a resolution would be at the cost of inducing multiple parallel substitutions for other mutations, and thus, we conclude that more than 100 codons in the genome that are not subject to an overt mechanistic bias underwent multiple parallel mutations in the course of SARS-CoV-2 evolution during the COVID-19 pandemic.

One immediate explanation of this observation is that these sites evolve under positive selection. The possible alternatives could be that these sites are mutational hotspots or that the appearance of multiple parallel mutations was caused by numerous recombination events (either real or artifacts caused by incorrect genome assembly from mixed infections) in the respective genomic regions. Contrary to what one would expect under the hotspot scenario, we found that codons with many synonymous substitutions tend to harbor few non-synonymous substitutions, and vice versa (Fig. S6 A). When a moving average with increasing window size was computed, only a weak positive correlation was observed between the numbers of synonymous and non-synonymous substitutions (Figs. S6 B&C, S7). Most sites in the virus genome are highly conserved, the sites with most substitutions tend to reside in conserved neighborhoods, and the local fraction of sites that harbor at least one mutation strongly correlates with the moving average (Fig. S8). Together, these observations indicate that SARS-CoV-2 genomes are subject to diverse site-specific and regional selection pressures but we did not detect regions of substantially elevated mutation or recombination in general agreement with other studies(28) despite the role recombination might have played in zoonosis(29–34).

### Positively selected sites in SARS-CoV-2 proteins

Given the widespread purifying selection affecting evolving SARS-CoV-2 genomes, substantially relaxed selection at any site is expected to permit multiple, parallel non-synonymous mutations to the same degree that any site harbors multiple, parallel synonymous mutations. Thus, seeking to identify sites subject to positive selection, we focused only on those non-synonymous substitutions that independently occurred more frequently than 90% of all synonymous substitutions excluding the mutagenic NCN context (see Methods). Most if not all sites in the SARS-CoV-2 genome that we found to harbor such frequent, parallel non-synonymous substitutions outside of the NCN context can be inferred to evolve under positive selection (Table S2, List 1). The positively selected residues form a co-occurrence network that likely reflects epistatic interactions (Fig. 1D and Table S3, see Methods), in which the central hubs are D614G in the spike (S) protein and two adjacent substitutions in the nucleocapsid (N) protein, R203K and G204R, the three most common positively selected mutations (Fig. 1C) (35). Fig. 1D,).

### Positively selected amino acid replacements in the receptor-binding domain of the spike protein

Spike D614G appears to boost the infectivity of the virus, possibly, by increasing the binding affinity between the spike protein and the cell surface receptor of SARS-CoV-2, ACE2(36). Conclusively demonstrating selection for a single site has proven challenging(37), even within this robust dataset. Although the emergence of this mutation corresponds to the extinction of partitions lacking 614G (see below), the possibility remains that this mutation is a passenger to some other mutagenic or epidemiological event. The 614 site of the S protein is evolutionarily labile, so that the ancestral reconstruction includes multiple gains of 614D after a previous loss. As a result, the reverse replacement G614D appears often enough to pass our statistical criteria for positive selection. Although severely biasing against recent events, one can additionally require that the mean tree fraction descendant from each candidate positively selected amino acid replacement be sufficiently large, removing from consideration events which are frequent but shallow (see Methods). The addition of this criterion results in a “shortlist” of 22 residues subject to the strongest selection (Table S2, List 2) that do not include 614D.

Additionally, apart from the selective advantage of a single replacement, it should be emphasized that D614G (but not G614D) is a central hub of the epistatic network (Fig. 1D). Conceivably, epistatic interactions with this residue can result in ensembles of mutations which substantially increase fitness. The ubiquitous epistasis throughout molecular evolution(38–41) suggests the possibility that many if not most mutations, which confer a substantial selective advantage, do so only within a broader epistatic context, not in isolation. By increasing the receptor affinity, D614G apparently opens up new adaptive routes for later steps in the viral lifecycle. The specific mechanisms of such hypothetical enhancement of virus reproduction remain to be investigated experimentally.

In addition to 614G, 31 spike mutations, most within the RBD, are signature mutations for divergent clades v1-3; emergent variants vAfrica or vOceania (see below); or established variants B.1.1.7, B.1.1.7_E484K, B.1.258_delta, B.1.351, B.1.429, P.1, or P.2(42–47) (Table S4, List 1). Three of these signature mutations pass the strict criteria for positive selection: S|N501Y, S|S477N, and S|V1176F, and S|N501Y makes the shortlist of the 22 strongest candidates. H69del/V70del are signature mutations for variant B.1.258_delta and have been previously observed to have rapidly emerged in an outbreak among minks(48, 49). A two amino acid deletion (in our alignment this deletion resolves to sites 68/69 due to many ambiguous characters in this neighborhood) appears multiple times independently throughout the tree and is present in approximately one third of the European sequences from January, 2021(Fig. S9, deletions are not shown in Fig. 1B, see below).

Two sites within the RBD, N331 and N343, have been shown to be important for the maintenance of infectivity(50). As could be expected, these amino acid residues are invariant. Four more substitutions in the RBD, among others, N234Q, L452R, A475V, and V483A, have been demonstrated to confer antibody resistance(50). N234Q, A475V, and V483A were never or rarely found in our alignment but L452R is a signature of variant B.1.429. Although not meeting our criteria for positive selection, it appeared multiple times across the tree, including within partition 1. Of greatest concern is perhaps N501Y. This amino acid replacement is a signature of variants B.1.1.7, B.1.1.7_E484K, B.1.351, P.1; divergent clade v2; and emergent variant vAfrica. N501Y is among the 22 strongest candidates for positive selection and has been demonstrated to escape neutralizing antibodies(51). N501T in the same site is of additional concern(52) and has also been observed in mink populations(53). Additionally, S|N439K, a signature mutation for variant B.1.258_DELTA that has been demonstrated to enable immune escape(54), is observed in a large portion of the tree.

The emergence of multiple mutations associated with immune evasion during a period of the pandemic when the majority of the global population had remained naïve is striking. Such adaptations are generally expected to emerge among host populations where many individuals have acquired immunity either through prior exposure or vaccination(55–58). Furthermore, this pattern of, most likely independent, emergence of persisting variants among both human and mink populations suggests the possibility that these mutations represent non-specific adaptations acquired shortly after zoonosis.

The factors underpinning the evolution of viral life history traits after zoonosis, especially virulence, remain poorly understood(59) but apparently result from selective pressures imposed by both epidemiological parameters (host behavior)(60, 61), which may be conserved across a variety of novel hosts, and specific properties of the host receptor. Whereas emergent mutations in the RBD of SARS-CoV-2 are, for obvious reasons, surveyed with great intensity, we have to emphasize the enrichment of positively selected residues in the N protein, which might relate to more deeply taxonomically conserved routes of host adaptation for beta-coronaviruses.

### Amino acid replacements associated with the nuclear localization signals in the nucleocapsid protein

Evolution of beta-coronaviruses with high case fatality rates including SARS-CoV-2 was accompanied by accumulation of positive charges in the N protein that might enhance its transport to the nucleus(62). Thirteen amino acid replacements in the N protein are signatures among the variants or major partitions discussed here, 7 of which: 203K, 204R, 205I, 206F, 220V, 234I, and 235F, are in the vicinity of the known NLS motifs or other regions responsible for nuclear shuttling(26). Two additional substitutions, 194L and 199L, rose to prominence in multiple regions during the summer of 2020. Two of these NLS-adjacent amino acid replacements, R(agg)203K(aaa) and G(gga)204R(cga), almost always appear together. This pair of substitutions includes the second and third most common positively selected sites after S614, and although another adjacent site, S(agt)202N(aat) is not a signature mutation, it is the 8^th^ most common positively selected residue. Among the 22 nonsynonymous substitutions that are apparently subject to the strongest selection (Table S2, List 2), 6 are in the N protein (202N, 203K, 204R, 234I, 292T, and 376T).

The replacements R(agg)203K(aaa) and G(gga)204R(cga) occur via three adjacent nucleotide substitutions. R(agg)203K(aaa) resolves to two independent mutations in the ancestral reconstruction: first, R(agg)203K(aag), then K(aag)203K(aaa). Furthermore, the rapid rise of 220V (excluded from consideration as a candidate for positive selection in our analysis due to its NCN context) in a European cohort during the summer of 2020 might be related to a transmission advantage of the variant carrying this substitution(63).These substitutions, in particular G(gga)204R(cga), which increases the positive charge, might contribute to the nuclear localization of the N protein as well. This highly unusual cluster of multiple signature and positively selected mutations across 5 adjacent residues in the N protein is a strong candidate for experimental study that could illuminate the evolution and perhaps the mechanisms of SARS-CoV-2 pathogenicity.

In addition to the many mutations of interest in the N and S proteins, Orf3a|Q57H is a signature mutation for partitions 6, 7, and v1. Q57H is the 4^th^ most common positively selected mutation. Although not considered a candidate for positive selection in our analysis due to its NCN context, ORF8 S84L is a hub in the larger epistatic network including all strongly associated residues (Fig. S10).

We also identified numerous nonsense mutations. Of particular interest seems to be ORF8|Q27*, which is a signature for variants B.1.1.7 and B.1.1.7_E484K and could be epistatically linked to positively selected residues including N|R203K and S|D614G. ORF8 has been implicated in the modulation of host immunity by SARS-CoV-2, so these truncations might play a role in immune evasion(64, 65).

### Potential role of epistasis in the evolution of SARS-CoV-2

Epistasis in RNA virus evolution, as demonstrated for influenza, can constrain the evolutionary landscape and promote compensatory variation in coupled sites, providing an adaptive advantage which would otherwise impose a prohibitive fitness cost(66–68). Because even sites subject to purifying selection can play an adaptive role through interactions with other residues in the epistatic network(69), the networks presented here (Figs. 1D, S10) likely underrepresent the extent of epistatic interactions occurring during SARS-CoV-2 evolution. The early evolutionary events that shaped the epistatic network likely laid the foundation for the diversification of the virus relevant to virulence, immune evasion, and transmission. As discussed above, these early mutations (including S|G614D) might provide only a modest selective advantage in isolation but exert a much greater effect through multiple epistatic interactions.

The epistatic network will continue to evolve through the entirety of the pandemic, and indeed, all emerging variants at the time of this writing are defined not by a single mutation but by an ensemble of signature mutations. Moreover, in addition to the apparent widespread intra-protein epistasis, there seem to exist multiple epistatic interactions between the N and S proteins. In particular, S|N501Y and N|S235F are both signature mutations for variants B.1.1.7 and B.1.1.7_E484K (Table S4, List 2) and this pair is in the top 25% of co-occurring pairs in our network ranked by lowest probability of random co-occurrence.

As with early founder mutations, when a new variant emerges with multiple signature mutations, it is unclear which, if any, confer a fitness advantage. Although it is natural to focus on substitutions within the RBD, we emphasize that all emergent variants contain substitutions in in the vicinity of known NLS motifs. In fact, the most statistically significant signature mutation (based on the Kullback-Leibler divergence) for vAfrica (consistent with variant B.1.351, see below) is N|T205I. As we suggest for S|D614G, these variant signature mutations are likely to exert a greater influence through multiple epistatic interactions than in isolation and each signature mutation can be a member of multiple epistatic ensembles beyond the group of signature mutations within which it was originally identified. Indeed signature mutations are shared among defined variants and we find evidence for an additional 18 putative epistatic interactions between variant signature mutations and other events throughout the tree which are not identified as signature mutations for any defined variant (Table S4, List 3). The growing ensemble of signature mutations that appear to be subject to positive selection and the existence of a robust network of putative epistatic interactions including these signatures, suggest that ongoing virus diversification is driven by host adaptation rather than occurring simply by neutral drift.

### Epidemiological Trends and Ongoing Diversification of SARS-CoV-2

Analysis of within-patient genetic diversity of SARS-CoV-2 has shown that the most common mutations are highly diverse within individuals(70–72). Such diversity could either result from multiple infections, or otherwise, could point to an even greater role of positive selection affecting a larger number of sites than inferred from our tree. Similarly to the case of Influenza, positive selection on these sites could drive virus diversification and might support a regular pattern of repeat epidemics, with grave implications for public health. An analysis of the relationships between the sequencing date and location of each isolate and its position within the tree can determine whether diversification is already apparent within the evolutionary history of SARS-CoV-2.

We first demonstrated a strong correlation between the sequencing date of SARS-CoV- 2 genomes and the distance to the tree root (Fig. S11), indicating a sufficiently low level of noise in the data for subsequent analyses. Examination of the global distribution of each of the major SARS-CoV-2 partitions (Figs. S12-14) indicates dramatic regional differences and distinct temporal dynamics (Fig. 2). A measure of virus diversity is necessary to map to these trends. We considered two modes of diversity. Intra-regional diversity reflects the mutational repertoire of the virus circulating in any individual region within any window of time. To measure intra-regional diversity, we sampled pairs of isolates from each region and timepoint and computed the mean tree-distance for a representative ensemble of these pairs. We found that intra-regional diversity has been steadily increasing throughout the entirety of the pandemic, with the exception of Oceania from June-August, 2020 (Figs. 3A/B) which corresponds to the period following a bottleneck in the total number of infections (Fig. S15) within that region. This unabated intra-regional diversification is a further evidence of a large repertoire of host-adaptive mutations of SARS-CoV-2 evolving within the human population.

**Figure 2.**
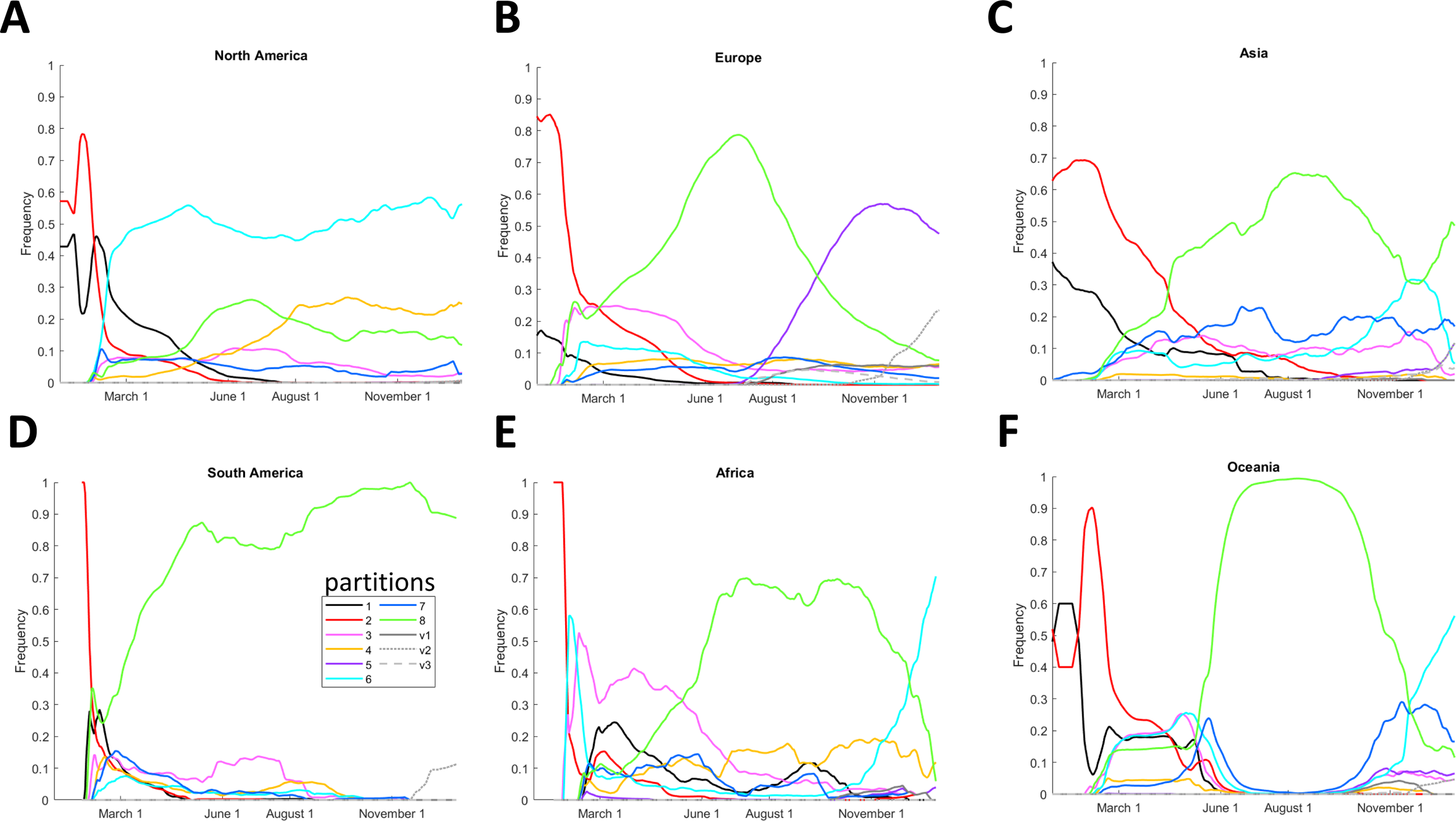
Regional SARS-CoV-2 partition dynamics during the COVID-19 pandemic. Probability distributions shown, for the absolute number of sequences, see Fig. S15.

The inter-regional diversity measures the degree to which the virus can be categorized into region-specific subtypes. A demonstration of substantial inter-regional diversity would perhaps constitute the most compelling and concerning evidence of the potential for repeat epidemics. We developed two measures of inter-regional diversity. The first one is analogous to the intra-regional diversity measure. We sampled pairs of isolates within each region and between each pair of regions within the same time window, computed the mean tree-distance for both representative ensembles of these pairs (intra- and inter-regional pairs), and calculated the ratio of inter-regional and intra-regional values (Fig. S16). The second one is a partition-level measure. For every pair of regions over each time window, we computed the Hellinger distance of the 11-group frequency distribution between all pairs of regions over each time window. (See Methods for details).

Both measures of inter-regional diversity support the division of the pandemic, through the beginning of 2021, into four periods (Fig1. 3C). The first period that ended in February 2020 represents rapid diversification into region-specific phylogenies. This period was followed by a major extinction event and global homogenization ending in March 2020. The following five months, March-July, represented a period of stasis, in terms of inter-regional diversity. Finally, July 2020 was the start of the ongoing period of inter-regional diversification.

The extinction of the earliest partitions, 1 and 2, corresponds to the advent of S|D614G, which became fixed in all other partitions and was globally ubiquitous by June 2020 (Fig. 3D). Partition 8, the only partition where N|203K and 204R were fixed, became dominant in every region outside of North America in the period that followed (Fig. S17). However, this did not result in a global selective sweep that would involve the extinction of partitions 1-7. Instead, multiple NLS-associated mutations rose to prominence across different partitions, becoming globally dominant by September (Figs. 3D, S18). To resolve this trend, at least two principal variants, N|203K/204R in partition 8 and N|220V in partition 5, have to be considered, and we identified 6 key amino acid replacements of interest for this period (N|203K/204R, N|220V, N|199L, N|194L, N205I, N206F).

**Figure 3.**
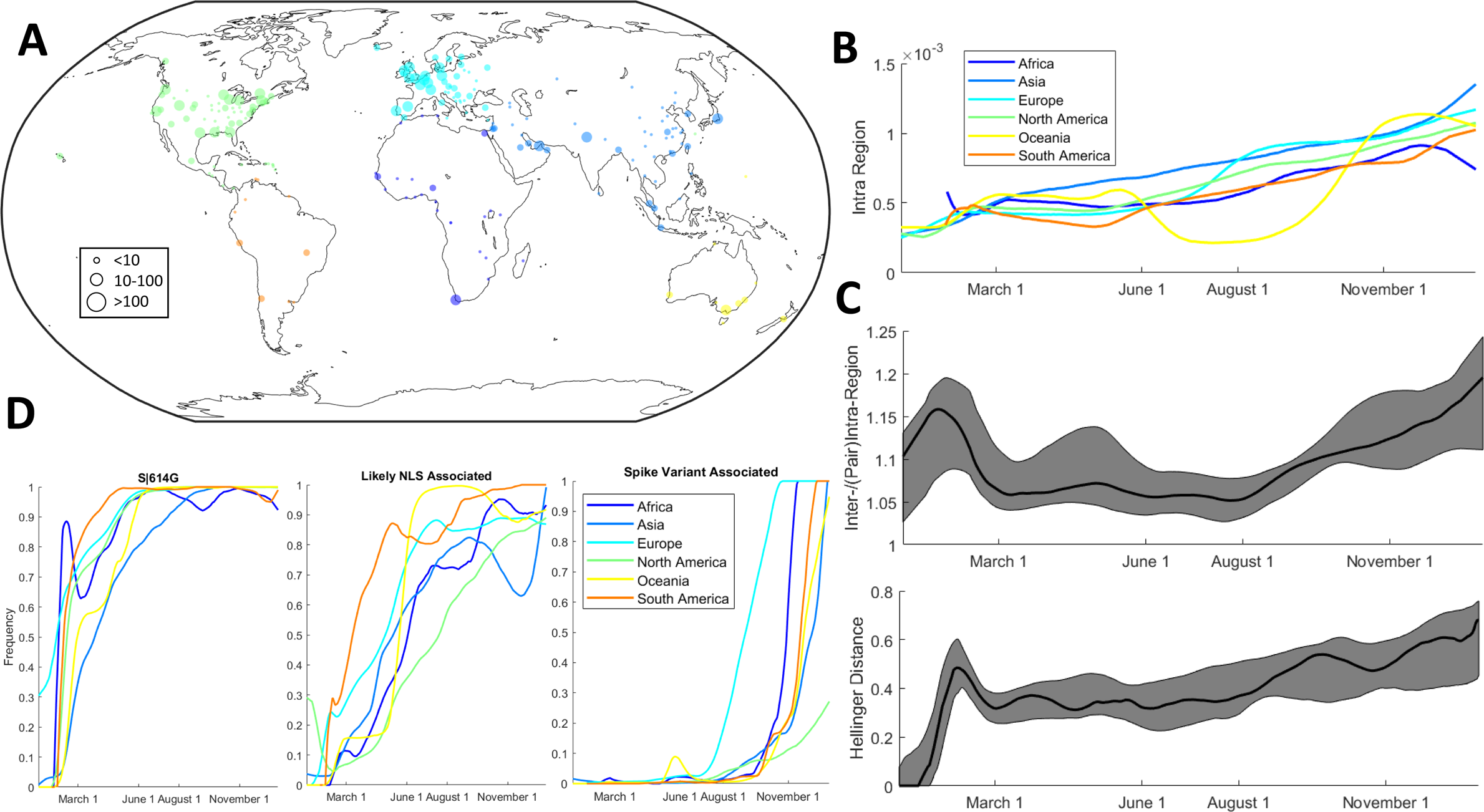
Global and regional trends in SARS-CoV-2 evolution. A. Global distribution of sequences with sequencing locations in each of the six regions considered. Color scheme is for visual distinction only. B. Intra-regional diversity measured by the mean tree-distance for pairs of isolates. C. (Top) The Hellinger distance for all pairs of regions over the 11 partition/clade distribution. 25th, 50th, 75th percentiles shown. (Bottom) The ratio of the mean tree-distance for pairs of isolates between regions vs. isolates within regions. 25th, 50th, 75th percentiles shown. D. The frequency of S|614G, at least one NLS-associated variant (N|194L, N119L, N203K, N205I, and N220V), and at least one emerging spike variant (Fig. S23, excluding S|477N).

In the next phase of the pandemic, partition 8 dramatically fell from dominance in two regions, Africa and Oceania, replaced by partitions 6 and 7. Although we did not find a distinct mutational signature associated with the rise of partition 7 in Oceania (Fig. S19), signatures associated with the rise of partition 6 were identified in both regions (Figs. S20-21). Neither of these two groups of sequences (late sequences from partition 6, Oceania and Africa, respectively) form topologically distinct clades; however, due to the conserved mutational signatures, we considered both groups to represent distinct emerging variants, vOceania and vAfrica. The signature for vAfrica is consistent with variant B.1.351. Additionally, two divergent clades within partition 8 and one clade within partition 3 emerged.

The most prominent is clade v2 with a signature consistent with variant B.1.1.7. Altogether, resolving this trend of emerging substitutions in the RBD (Figs. 3D, S22-24) requires the consideration of at least 3 variants and includes 59 signature mutations.

Clade v1 appeared first in Europe in April, 2020, v2, also in Europe, in September, 2020, and V3 in Asia and North America, in April, 2020 (Fig. S25). Also notably, although S|477N initially appears in February/March, 2020 in Europe, Oceania, and North America, it dramatically rises to prominence in Oceania in April, about 3 months before this mutation becomes prominent elsewhere. S|477N is a signature mutation for v1 stemming from partition 3; however, the sequences from Oceania bearing this mutation from Summer, 2020 are in partition 8. The dramatic diversity of signature mutations among these variants decreases the likelihood of future selective sweeps (in the absence of bottlenecks in the total number of infected hosts) and increases the likelihood of repeat epidemics.

### The impact of SARS-CoV-2 Diversification on Testing and Vaccination

The ongoing diversification of SARS-CoV2 poses problems for both testing and vaccination. Substitutions in the E protein have already been demonstrated to interfere with a common PCR assay(73). Generally, ORF1ab is more conserved than the S protein, which itself is more conserved than the remaining ORFs (Figs. S2-3). Using our SARS-CoV-2 MSA, we surveyed 10 regions from ORF1ab(5), N(4), and E(1) genes that are commonly used in PCR assays(74) for substitutions relative to the reference sequence. Among the more than 175k genome sequences, there were thousands of nucleotide substitutions in each of these regions, but those in ORF1ab were markedly less variable than those in N (Supplementary table 5), with one region in N demonstrating variability in nearly one third of all isolates. It can be expected that most targets within the polyprotein will remain subject to the fewest polymorphism-induced false negatives even as the virus continues to diversify.

Of the 9 primary vaccines/candidates (75), three are inactivated whole-virus (Sinovac, Wuhan Institute of Biological Products/Sinopharm, Beijing Institute of Biological Products/Sinopharm); five utilize the entire spike protein as the antigen (Moderna/NIAID, CanSino Biological Inc./Beijing Institute of Biotechnology, University of Oxford/AstraZeneca, Gamaleya Research Institute, Janssen Pharmaceutical Companies) and one utilizes only the RBD (Pfizer/Fosun Pharma/BioNTech). In addition to the greater sequence conservation of the spike protein relative to all other ORFs outside of the polyprotein, it is the principal host-interacting protein of SARS-CoV-2, making both the whole protein and the RBD obvious antigenic candidates. Most mutations in the RBD were demonstrated to decrease infectivity, but some conferred resistance to neutralizing antibodies(49). Multiple mutations in the RBD are signature mutations in emerging variants and some have been demonstrated to result in neutralizing antibody evasion(51). Different choices of the antigen could result in more or less generalizable immunity to these variants.

## Conclusions

Virus evolution during a pandemic is a fast moving target, and unavoidably, aspects of this analysis will be outdated by the time of publication. Nevertheless, several trends revealed here appear general and robust. Although it is difficult to ascertain positive selection for individual sites, the overall adaptive character of SARS-CoV-2 evolution involving multiple amino acid replacements appears to be beyond reasonable doubt. As expected, there are multiple positively selected sites in the S protein, but more surprisingly, N protein includes several sites that appear to be strongly selected as well. The involvement of these adaptive substitutions in the nuclear localization of the N protein appears likely. Importantly, some of the mutations, for which positive selection was inferred, co-occur on multiple occasions and seem to form a robust epistatic network. Most likely, the effect of positive selection is manifested primarily at the level of epistatic interactions.

Clearly, despite the dramatic reduction of global travel(76), the evolution of SARS-Cov- 2 is partly shaped by globalizing factors, including the increased virus fitness conferred by S|D614G, N|R203K&G204R, and other positively selected substitutions. However, we obtained strong evidence of both continuous virus diversification within geographic regions and “speciation”, that is, formation of stable, diverging region-specific variants. This ongoing adaptive diversification could substantially prolong the pandemic and the vaccination campaign, in which variant-specific vaccines are likely to be required.

## Author contributions

EVK initiated the project; NDR and GF collected data; NDR, GF, YIW, PM, FZ and EVK analyzed data; NDR and EVK wrote the manuscript that was edited and approved by all authors.

## Acknowledgements

The authors thank Koonin group members for helpful discussions. NDR, YIW PM, and EVK are supported by the Intramural Research Program of the National Institutes of Health (National Library of Medicine).

## Methods

### Multiple alignment of SARS-CoV-2 genomes

All available SARS-CoV-2 genomes as of January 8, 2021 were retrieved from the Genbank(13), Gisaid(14), and CNCB(15) datasets. Sequences with apparent anomalies (sequence inversion etc.) were immediately discarded. Sequences were harmonized to DNA (e.g. U was transformed to T to amend software compatibility) and clustered according to 100% identity with no coverage threshold using CD-HIT(77, 78), with ambiguous characters masked. All characters excepting ACGT were considered ambiguous. The least ambiguous sequence from each cluster was selected and sequences shorter than 25120 nucleotides were discarded.

Exterior ambiguous characters (preceding/succeeding the first/last defined nucleotide) were removed, and sequences with more than 10 remaining interior, ambiguous characters were discarded. A reference alignment was previously constructed using the same protocol as follows with the exception of the --keeplength specification in November, 2020. The updated database was aligned using multi-threaded MAFFT(79) with 80 cores (--thread 80, when more cores were allocated they were not utilized) and 3.8Tb of RAM to maintain usage of the normal DP algorithm(79) (--nomemsave) against this reference alignment (specifying --keeplength). Aligning “from scratch” without -- keeplength proved to be prohibitively slow so we recommend first constructing a reference alignment from a suitable subset of sequences. Sequences sourced from non-human hosts were manually identified from the metadata and those excluded at the previous step were added to the alignment using MAFFT, (again specifying -- keeplength). Note that use of the --keeplength option will not include insertions relative to the reference alignment.

Sites corresponding to protein-coding ORFs were then mapped to the alignment from the reference sequence NC_045512.2 excluding stop codons as follows: 266- 13468+13468-21552, orf1ab; 21563-25381, S; 25393-26217, orf3a; 26245-26469, E; 26523-27188, M; 27202-27384, orf6; 27394-27756, orf7a; 27756-27884, orf7b; 27894-28256, orf8; and 28274-29530, N. The remaining sites were discarded.

The resulting alignment contained out-of-frame gaps. Gaps in the reference sequence, corresponding to insertions, were found to correspond to gaps in all but fewer than 1% of the remaining sequences (all gaps in the reference sequence correspond to gaps in the alignment from November, 2020, the use of --keeplength prohibited the recognition of any insertions relative to the reference sequence which were not present in this reference alignment). These sites were discarded. The remaining gaps, corresponding to deletions relative to the reference sequence, shorter than three nucleotides were replaced with the ambiguous character, N. Longer gaps were shifted into frame and padded with ambiguous characters on either end of the gap, minimizing the number of sites altered.

A fast, approximate tree was then built using FastTree(80) (parameters: -nt -gtr -gamma -nosupport -fastest) to unambiguously define two clusters of sequences: an outgroup consisting of 14 sequences sourced from non-human hosts prior to 2020 and the main group. The tree construction requires the resolution of very short branch lengths which makes it necessary to compile FastTree at double precision. Outliers from the remaining sequences were then identified based on the Hamming distance (excluding gaps and ambiguous characters) to the nearest neighbor, the Hamming distance to the consensus, and the degree to which those substitutions relative to consensus were clustered in the genome. At this step, 81 sequences were removed.

The resulting alignment, consisting of 98,185 sequences and 29,119 sites, was maintained for the construction of the global tree and ancestral sequence reconstruction. In an effort to minimize the impact of sequencing error on the tree topology, as well as to decrease computational costs, a reduced alignment was then constructed through the removal of 1) invariant sites, 2) sites invariant with the exception of a single sequence, and 3) sites invariant throughout the main group with the exception of at most one sequence representing each minority nucleotide.

Removing these sites created substantial redundancy, so a representative sequence was selected for each cluster of 100% identity to yield an alignment consisting of 90,585 sequences and 16,487 sites. As described below and in the main text, a third alignment was constructed including only the top 5% of sites with the most common substitutions relative to consensus (of this second alignment) and again removing redundant sequences to yield 32,563 sequences and 834 sites.

### Tree Construction

We sought to optimize tree topology with IQ-TREE(81); however, building the global tree was computationally prohibitive, and thus, we proceeded to subsample the smallest alignment (834 sites) as follows. First, a core set of maximally diverse sequences is selected. The set is initialized with a pair of sequences: a sequence maximizing the number of substitutions relative to consensus and a paired sequence which maximizes the Hamming distance to itself. Sequences are then added to this core set one at a time maximizing the minimum Hamming distance to any representative of the set until *N* sequences are incorporated. Next, *ceil(L*/(*M*−*N*)) resulting sets are initialized with this core set where *M* is the target number of sequences and *L* is the total number of sequences in the alignment (32, 363). Then, sequences that have not yet been incorporated into any resulting set are added to each resulting set, again one at a time, maximizing the minimum distance to any representative of the set until *M* sequences are incorporated. The order of the resulting sets is randomized at each iteration without repeats. Once every (main group) sequence has been incorporated into at least one resulting set, sequences are randomly incorporated into each set until every set contains *M* sequences. Finally, the outgroup is added to each resulting set. We chose *M*=3,000 in an effort to optimize computational efficiency and *N*=300. Note that while increasing *N* increases the number of sets required for alignment coverage, and thus compute time, insufficient overlap between the sequences assigned each sub-alignment greatly affects the results of subsequent steps. As discussed in the main text, executing this protocol on an alignment containing most or all sites may not yield a consistent deep tree topology or “skeleton” since maximizing the hamming distance of any subset over all sites does not guarantee maximizing the tree distance in the resultant global topology. This is why limiting the alignment to sites with common substitutions relative to consensus is essential at this step.

A tree was then built, using IQ-TREE, for each maximally diverse set, with the evolutionary model fixed to GTR+F+G4 and the minimum branch length decreased from the default 10e-6 to 10e-7, according to the results of previous parameter studies(17).

These trees were then converted into constraint files and merged to generate a single global constraint file for use within FastTree (parameters: -nt -gtr -gamma -cat 4 - nosupport -constraints).

The remaining sequences excluded from this tree but present in the second alignment (90,585 sequences and 16,487 sites) were then reintroduced as unresolved multifurcations and a new constraint file from the multifurcated tree was constructed. A second iteration of FastTree was initiated on the second alignment to produce an intermediate tree. This tree was primarily constructed as an intermediate step to limit the impact of sequencing errors on the final topology as mentioned in the main text; however, it is also less computationally intensive. The last step was then repeated on this intermediate tree to construct the global topology for the whole alignment. The final, global tree was rooted at the outgroup.

### Reconstruction of Ancestral Genome Sequences

Ancestral states were estimated by Fitch Traceback(18). Briefly, character sets were constructed from leaf to root where each node was assigned the intersection of the descendant character sets if not empty and the union otherwise. Then, moving from root to leaf, nodes with more than one character in their set were assigned the consensus character if present in their set or a randomly chosen representative character otherwise. Substitutions between states were identified and placed in the middle of the branch bridging the pair of nodes.

Statistical associations between mutations were computed in a manner similar to that previously described(35). Briefly, sequences were leaf-weighted based on the branch lengths of the ultrameterized, tree. Every mutation present across the tree at 200 mean leaf-weight equivalents or more was considered. The probability of independent co-occurrence between any pair was estimated in two ways. An arbitrary member of the pair was selected as the ancestral mutation, and the binomial probability:

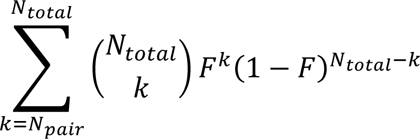

was computed where *N_total* is the number of substitutions to the descendant mutation across the entire ancestral record, *N_pair* is the number of substitutions to the descendant which succeed or appear simultaneously with a substitution to the ancestral mutation, and *F* is the fraction of the tree (fraction of all applicable branch lengths) occupied by the ancestral mutation. The ancestral/descendent designation was then reversed and the “binomial score” was constructed as the negative log of the product of these two terms. Additionally, for each pair, the observed and expected (product of the tree fractions) tree intersections were calculated and the “Poisson score” (analogous to the log-odds ratio) was calculated:

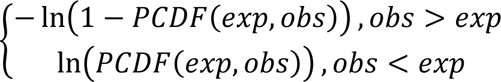

where PCDF(exp,obs) is the cumulative probability of a Poisson distribution with mean “exp”, the expected value of the data, and evaluated at “obs”, the observed value of the data. Both scores are reported. Table S3 displays putative positively selected mutations with both scores above 5 or at least two simultaneous substitutions. Fig. 1D only displays associations between mutations in the N or S proteins. Fig. S10 does not exclude mutations with NCN context but meets all other statistical criteria for positive selection and does not display mutations in the polyprotein.

### Classical Multidimensional Scaling of the MSA

Pairwise Hamming distances were computed for all pairs of rows in the global MSA ignoring gaps and ambiguous characters i.e. the sequences *X*=”ATN-A” and *Y*=”NTAAT” would be assigned a distance of 1. The resulting distance matrix was embedded in three dimensions with the MATLAB(82) routine “cmdscale”. 100 rounds of stochastically initiated k-means clustering of the embedding was conducted and the optimum cluster number was determined to be 5 on the basis of the silhouette score distribution (Fig S1).

### Validation of Mutagenic Contexts

Mutations were divided into four categories: synonymous vs non-synonymous substitutions and high vs low frequency of independent occurrence. For example, consider codon X with 3 non-synonymous substitutions gat->ggt and 1 non-synonymous substitution gat->cgt. In this context, a non-synonymous nucleotide substitution a->g of frequency 4 would be recorded in nucleotide (X-1)*3+2. The low vs high frequency threshold was determined by the 90^th^ percentile of the synonymous mutation frequency distribution (operationally 7). For each mutation, the trinucleotide contexts from the ancestral reconstruction at the nodes where the mutation occurred were compared to the background genome-wide frequencies, computed for the inferred common ancestor of SARS-CoV-2.

The expected frequencies of the trinucleotides using the background distribution were tabulated; the Yates correction (+/-0.5 to the original count depending on whether the count is below or above the expectation) was applied to the observed frequencies; the log-odds ratios of the (corrected) observed frequencies to the expectation were computed; and CMDS was applied to the Euclidean distances between the log-odds vectors to embed the points onto a plane (Fig. S4 A.). This analysis was then repeated, this time, distinguishing only between high and low frequency substitutions but not N and S (Fig. S4 B). Finally, the differences in the contexts of high frequency synonymous vs non-synonymous events were considered in the same manner and the chi-square statistics ((observed-expected)^2/expected) were compared with the critical chi-square value (p=0.05/64, df=1, Fig. S4 C.).

### Computation of *dN/dS*

For each of the 24 ORFs (splitting orf1ab into 15 segments corresponding to the 15 mature proteins, nsp11 and nsp12 combined), 10 reduced alignments were constructed as follows. Sequences were ordered based on diversity, in the same order with which they were included in the constraint trees. The first 10 sequences are conserved across every alignment and the remaining 40 are unique to each alignment. The reference sequence, NC_045512.2, was additionally added to each reduced alignment. PAML(83) was then used to estimate tN, tS, *dN/dS*, N, S, and N/S for each segment and every reduced alignment.

Given the global ancestral reconstruction from Fitch traceback, the total number of non-synonymous and synonymous substitutions (nN and nS, respectively) as well as these tallies normalized by the respective segment length (tN, and tS, respectively) were retrieved for each segment. . A hybrid *dN/dS* value for each segment was estimated to be (nN/nS)/(N/S)* where (N/S)* is the median value of N/S across all repeats for the segment.

### Metadata Assignment

Headers for all isolates belonging to CD-HIT clusters with a representative incorporated into the alignment with fewer than 10 interior ambiguous characters were processed to extract the sequencing date and location. Sequencing location abbreviations were matched to full names and the latitude/longitude of a representative city for each location was retrieved from simplemaps (https://simplemaps.com/data/world-cities)(84).

### Regional Divergence Analysis

Two approaches, one partition dependent and one partition independent, were used as described in the main text. The Hellinger distance between regions over a sliding time window was computed between regions for the 11 (partitions/variant clades) group distribution. Next, 400 isolates were randomly selected from each region over a sliding window and 200 pairs within each region as well as 200 pairs between each pair of regions were composed. The tree distance between each pair was computed and the mean for each inter- and intra-regional pair tree-distance distribution was recorded. In Figs. 3C and S16, the 25^th^, 50^th^, and 75^th^ percentiles are shown of the 15 possible pairs of (6) regions. Regions are selected based on GISAID metadata. The inter-regional tree divergence (Figs. 3C, top and S16C) is reported as the ratio between the mean of the inter-regional pair tree-distance and the mean of the intra-regional pair tree distances across both regions.

## Supplemental Figures

Figure S1. 25^th^, median (solid line), and 75^th^ percentiles of the silhouette score distribution for 100 stochastically initiated rounds of k-means clustering for 2-16 clusters and a projection of the 3D embedding of the pairwise Hamming distance matrix between SARC-CoV-2 genomes. Partitions are color-coded and wires enclose the convex hulls for each of the five optimal clusters.

Figure S2. **A.** Distributions of the moving average, respecting segment boundaries, across a 100 codon window for synonymous (blue) and amino acid (orange) substitutions. Solid lines: normal approximations of the distributions (same median and interquartile distance); solid lines: approximation with the same median and theoretical (Poisson) variance. **B.** Moving averages, respecting segment boundaries, across a 100 codon window for synonymous and nonsynonymous substitutions per site, raw (top) and normalized by the median (bottom). There are several regions in the genome with an apparent dramatic excess of synonymous substitutions: 5’ end of orf1ab gene; most of the M gene; 3’-half of the N gene, as well as amino acid substitutions: most of the orf3a gene; most of the orf7a gene; most of the orf8 gene; and several regions in of the N gene.

Figure S3. Moving average over a window of 1000 codons, not respecting segment boundaries, of the total number of nucleotide exchanges n1->n2 summed over all substitutions. The ratio to the median over the entire alignment is also displayed as well as the normalized exchange distribution *(i.e.* #c->t/(#c->t+#c->g+#c->a)).

Figure S4 **A.** Two dimensional embedding of the Euclidean distances between the log-odds vectors of low and high frequency, nonsynonymous and synonymous mutations in the space of trinucleotide contexts relative to background expectation. The context of the high-frequency events (both S and N) is dramatically different from the background frequencies. There is a strong common component in the deviation of both kinds of high-frequency events. The context of the low-frequency events (both S and N) also differs slightly, in the same direction, from the background frequencies. There is a consistent distinction between synonymous and non-synonymous events, suggesting that a single mutagenic context or mechanistic bias does not account for both S and N events. **B.** Log odds ratio of low and high frequency mutations, both synonymous and nonsynonymous, relative to background expectation for each trinucleotide context. The NCN context (i.e. all mutations C->D) harbors dramatically more mutation events than the other contexts (all 16 NCN events are within the top 20 most-biased high-frequency events). The log-odds ratios for low-frequency events are poorly correlated with those for high-frequency events, suggesting that different mechanisms may be responsible for the strong bias observed among high frequency events and the weaker bias observed among low frequency events. **C.** Log odds ratio of high frequency nonsynonymous mutations relative to the background expectation from the sum of both high synonymous and high nonsynonymous mutations vs. the sum + 1. There are 20 contexts where synonymous and non-synonymous events differ significantly (chi-sq> 11.28). 2/9 contexts with an excess of non-synonymous events are NCN (gct,tct). The remaining 7 are NGN (agt,gga,aga,ggt,agc,tgt). This additionally suggests that these non-synonymous events could be driven by other mechanisms. There is no correlation between the frequency of event context and the log-odds ratio for non-synonymous events, further suggesting that the log-odds ratio is not biased by hot-spot mutation context.

Figure S5. Correspondence between the “tree length for dN”, “tree length for dS”, and *dN/dS* between PAML and the results of the ancestral reconstruction utilizing Fitch traceback across 24 ORFs. Three high outliers in the PAML tS distribution are identified in the third plot and omitted from the first two.

Figure S6. **A.** The number of nonsynonymous events vs the number of synonymous events per codon. **B.** The moving average of 100 codons, respecting segment boundaries. **C.** The moving average after removing outlier high frequency events. Rho refers to Spearman. Dashed lines are 2/1.3*x reflecting the genome-wide ratio of nonsynonymous to synonymous substitutions, solid lines are linear best fit. Red points correspond to the middle third of the N protein.

Figure S7. Moving averages across a 100 codon window for synonymous and nonsynonymous substitutions per site in the N protein after removing outlier high frequency events. The nonsynonymous substitution frequencies in the center of the protein are not elevated relative to either terminus.

Figure S8. The fraction of sites with at least one substitution vs moving averages, respecting segment boundaries, over windows of 100 codons for synonymous and nonsynonymous substitutions.

Figure S9. Site history trees for spike 69 as drawn in Fig. 1C.

Figure S10. Epistatic network for the tree including mutations with NCN context and meeting all other criteria for positive selection. Mutations in the polyprotein are not displayed.

Figures S11. Correlation between sequencing date and tree distance to the root for all isolates with metadata as well as those which appear explicitly in the tree.

Figures S12-14. Global distribution of sequences. Color represents the number of sequences from that location and size represents the fraction of sequences from the clade displayed. Partition indices are in the top left corner of each map.

Figure S15. Regional SARS-CoV-2 partition dynamics during the COVID-19 pandemic (absolute number of sequences shown in contrast to Fig. 2).

Figure S16. The mean tree distance between pairs of isolates **A.** from different regions, **A.** within the same region (averaged over both regions in each pair) and **C.** The ratio over time (see Methods). 25^th^, 50^th^, and 75^th^ percentiles of all 15 pairs of 6 regions. The ratio reported is between the mean of the inter-regional pair tree-distance and the mean of the intra-regional pair tree distances across both regions for each pair of regions.

Figure S17. Regional distributions of major partitions in the global topology March vs. July and July vs. November.

Figure S18. The frequencies of NLS-associated mutations N|194L, N119L, N203K, N205I, and N220V over time and across geographic regions along with S|614G for reference.

Figure S19. The Kullback-Leibler divergence and sequence logo for the 15 most divergent codons in sequences sourced after October 15, 2020 from Oceania in partition 7 vs. all sequences from Oceania in partition 7.

Figure S20. The Kullback-Leibler divergence and sequence logo for the 15 most divergent codons in sequences sourced after November 1, 2020 from Oceania in partition 6 vs. all sequences from Oceania in partition 6.

Figure S21. The Kullback-Leibler divergence and sequence logo for the 15 most divergent codons in sequences sourced after November 1, 2020 from Africa in partition 6 vs. all sequences from Africa in partition 6.

Figures S22-24. The frequencies of variant-associated mutations in the spike protein over time and geographic regions.

Figure S25. Regional SARS-CoV-2 variant clade dynamics during the COVID-19 pandemic (log of absolute number of sequences shown).

## Supplemental Tables

Table S1. The list of all mutations either in the top 100 most commonly observed or top 100 with the greatest number of parallel substitutions ordered as they appear in the genome.

Table S2. List of sites most likely to be evolving under positive selection. For List 2 the average tree fraction descendant from each candidate positively selected amino acid replacement must be sufficiently large(see Methods).

Table S3. All epistatic interactions among states meeting the criteria outlined in the main text for likely positive selection with binomial/Poisson scores greater than 5 or at least 2 simultaneous substitutions. Each mutation must have a minimum weight of approximately 200 leaves and each pair, 100 leaves. Each pair is arbitrarily ordered and the numbers of simultaneous, descendant, and independent substitutions are tabulated.

Table S4. **List 1**. List of variant mutations and variant IDs sorted by the number of variant ID’s assigned to each mutation. **List 2**. List of all pairs of mutations associated with a single variant ID (internal variant ID’s excluded. **List 3**. List of putative epistatic interactions between variant mutations and other states in the tree.

Table S5. The number of isolates (out of approximately 175k) observed to bear at least one substitution relative to the reference sequence, NC_045512.2, within the regions specified. These regions are commonly used within PCR assays for diagnostic testing.

## Notes

### Competing Interest Statement

The authors have declared no competing interest.

### Summary of Updates

This version of the manuscript has been revised to update the collection of the SARS-CoV-2 genomes that were analyzed to identify the evolutionary regime of the virus along with the global and regional evolutionary trends.

