## Supplemental figures for "Ongoing Global and Regional Adaptive Evolution of SARS-CoV-2"

**S1**

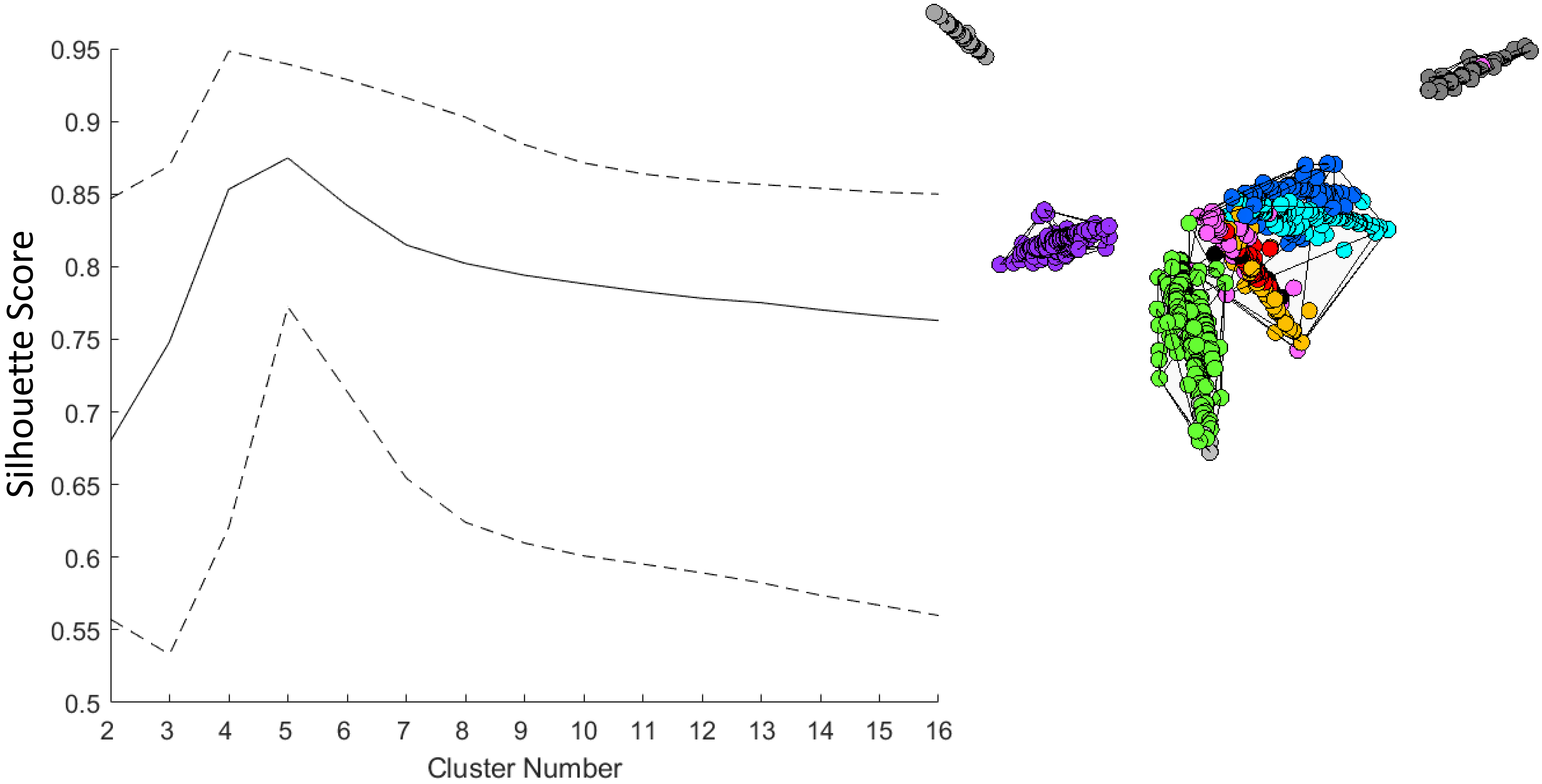
Silhouette Score
0.95
0.9
0.85
0.8
0.75
0.7
0.65
0.6
0.55
0.5
2 3 4 5 6 7 8 9 10 11 12 13 14 15 16
Cluster Number

# S2

**A**

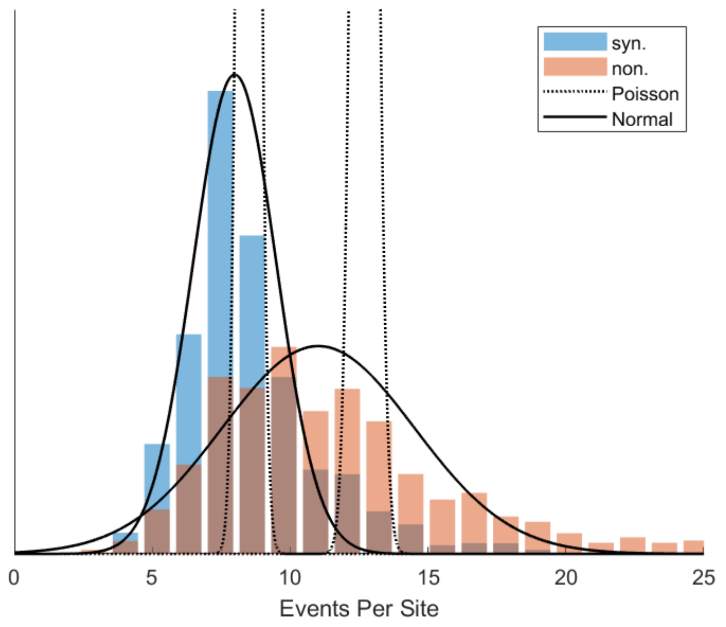
syn.
non.
Poisson
Normal
0
5
10
15
20
25
Events Per Site

**B**

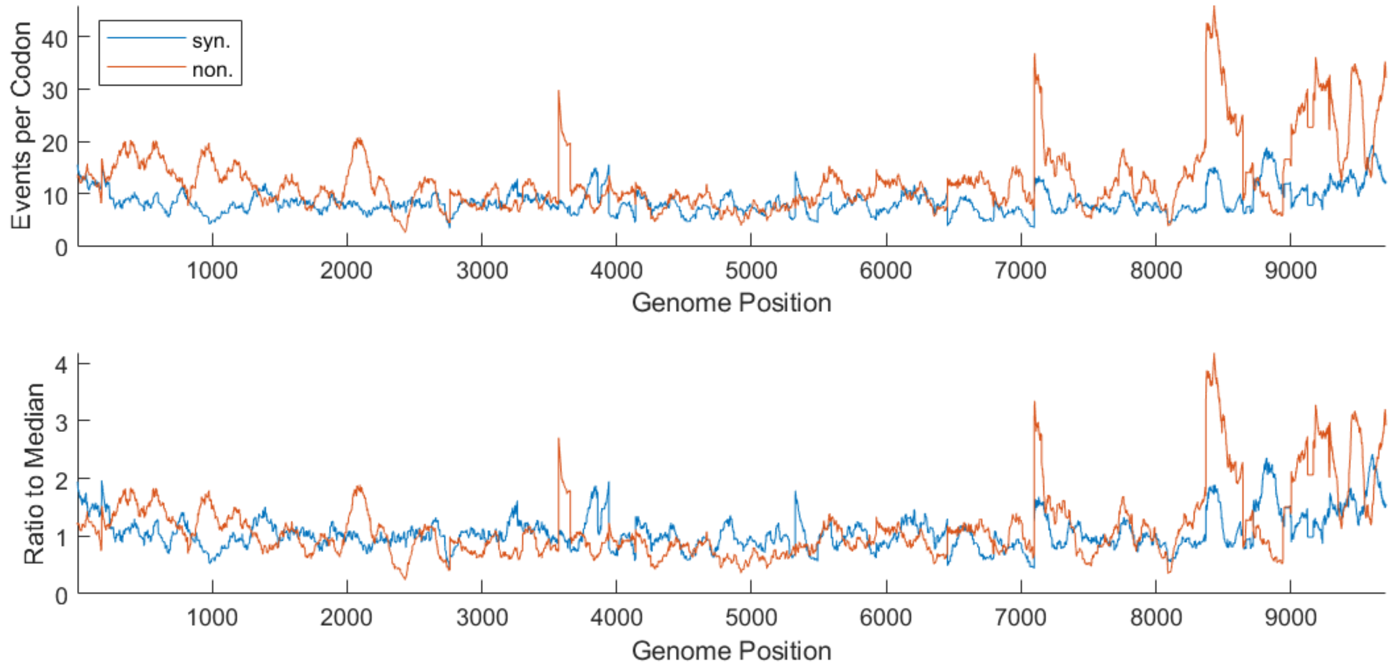
Events per Codon
syn.
non.
40
30
20
10
0
1000
2000
3000
4000
5000
6000
7000
8000
9000
Genome Position
Ratio to Median
4
3
2
1
0
1000
2000
3000
4000
5000
6000
7000
8000
9000
Genome Position

# S3

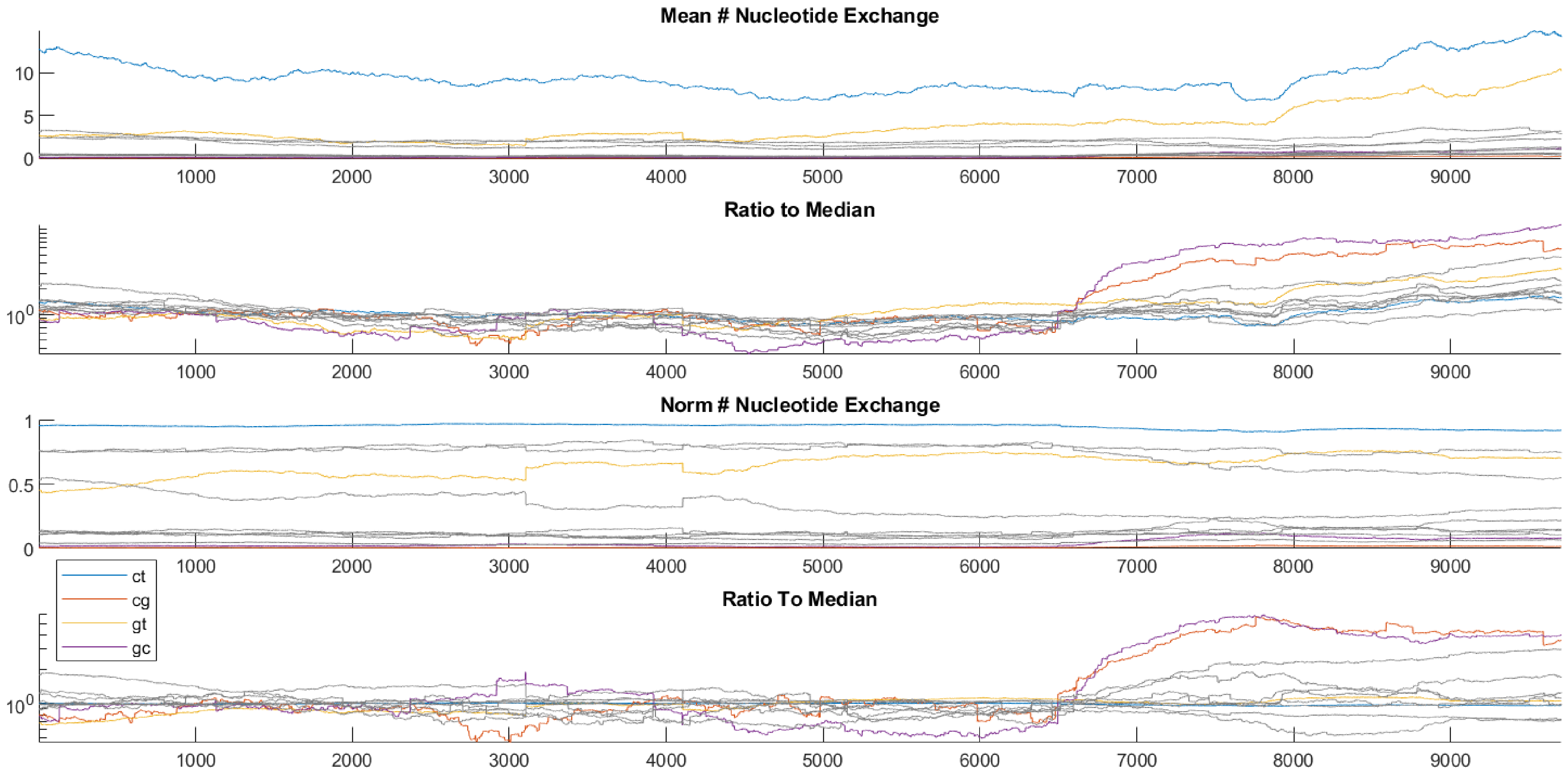
Mean # Nucleotide Exchange
10
5
0
1000
2000
3000
4000
5000
6000
7000
8000
9000
Ratio to Median
$10^0$
Norm # Nucleotide Exchange
1
0.5
0
ct
cg
gt
gc
Ratio To Median
$10^0$

**S4**

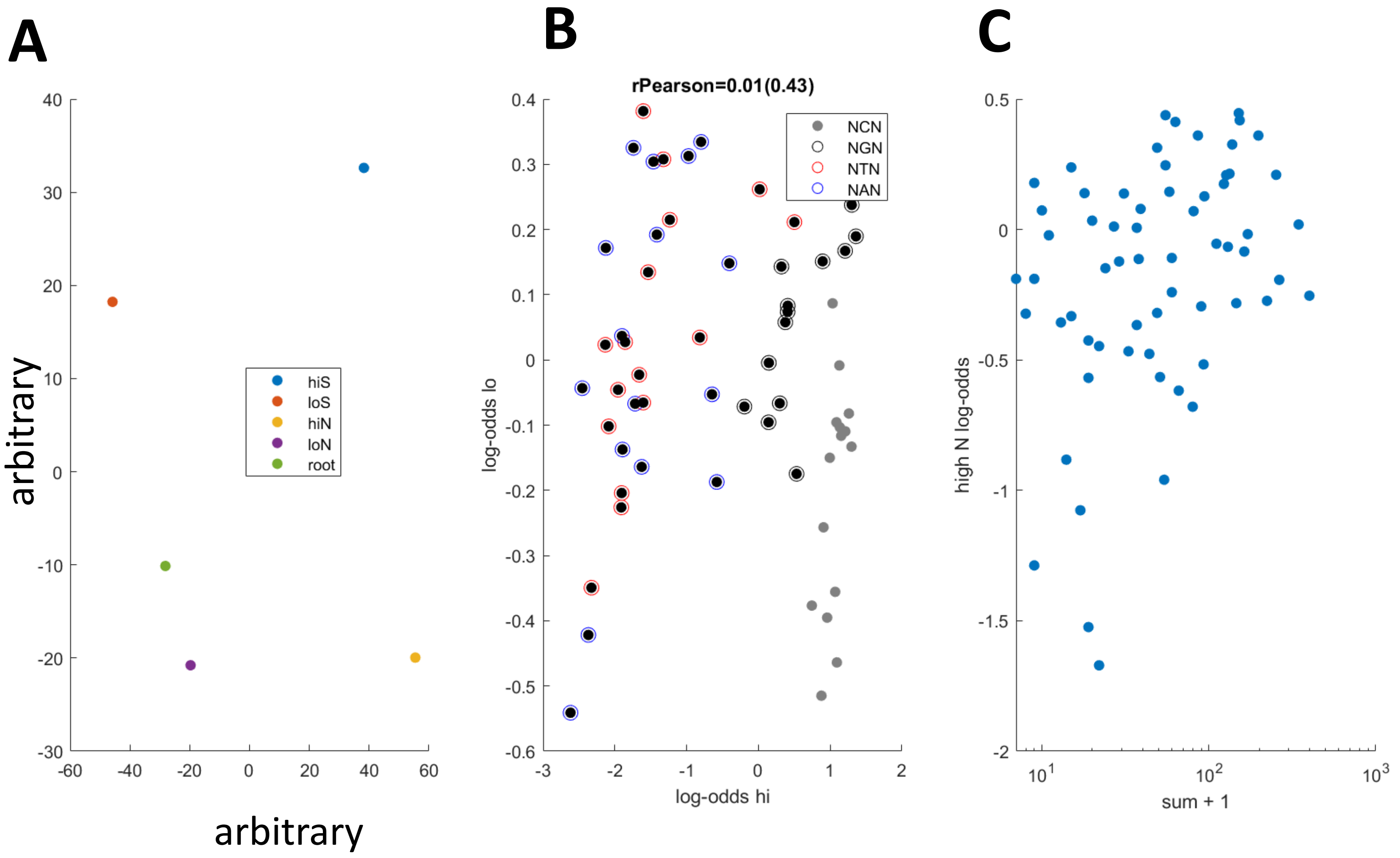


| context | log-odd S | log-odd N | chi-sq |
|---|---|---|---|
| agt | -2.04 | 0.45 | 72.63 |
| gga | -2.13 | 0.44 | 26.41 |
| aga | -1.63 | 0.42 | 62.85 |
| ggt | -1.68 | 0.41 | 25.9 |
| tgt | -1.09 | 0.36 | 55.91 |
| agc | -1.14 | 0.36 | 25.07 |
| tgc | -0.9 | 0.33 | 30.98 |
| gct | -0.42 | 0.21 | 19.83 |
| tct | 0.26 | -0.19 | 14.04 |
| aca | 0.32 | -0.25 | 34.32 |
| tca | 0.34 | -0.27 | 21.31 |
| cca | 0.34 | -0.28 | 14.46 |
| atc | 0.5 | -0.52 | 23.85 |
| gcg | 0.51 | -0.57 | 14.36 |
| tcg | 0.54 | -0.62 | 21.6 |
| ata | 0.58 | -0.68 | 30.33 |
| atg | 0.67 | -0.96 | 31.4 |
| cta | 0.76 | -1.52 | 16.31 |
| ttg | 0.79 | -1.67 | 20.8 |

# S5

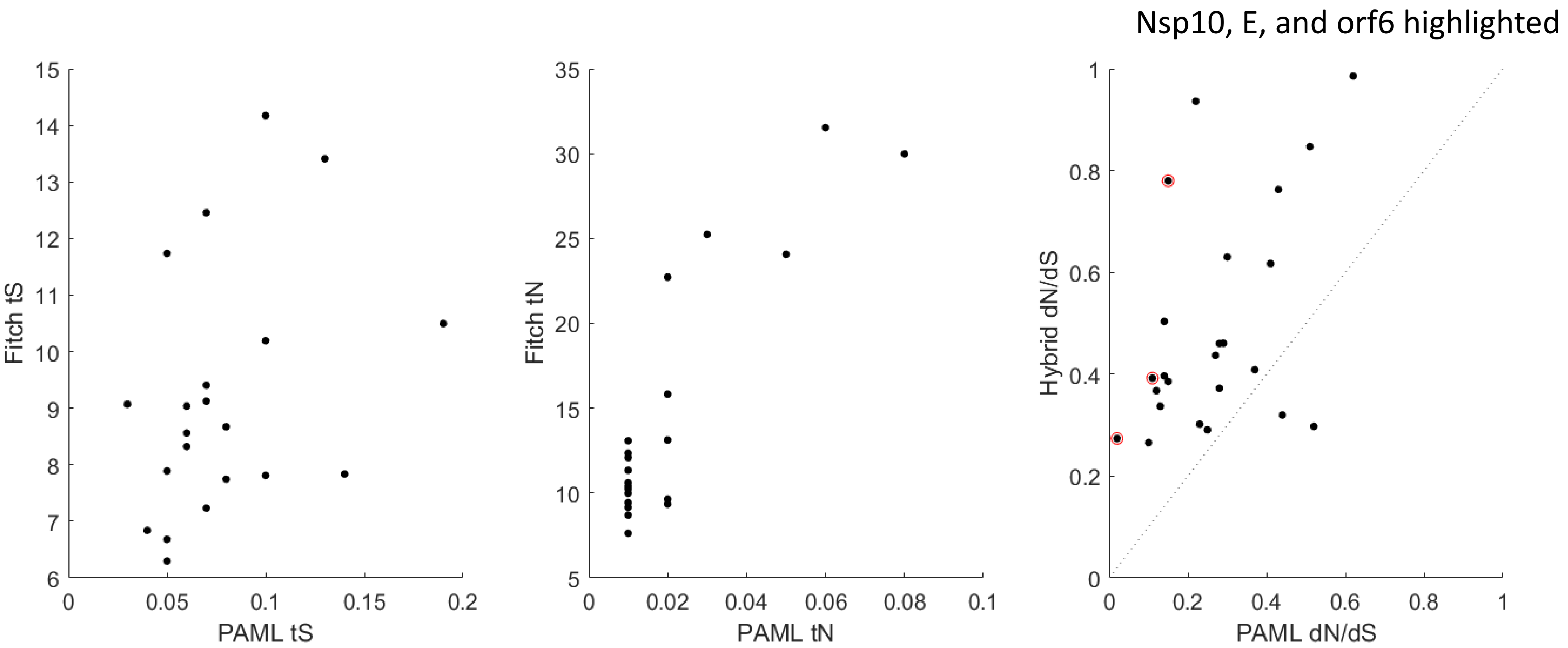

Nsp10, E, and orf6 highlighted
Fitch tS
PAML tS
Fitch tN
PAML tN
Hybrid dN/dS
PAML dN/dS
15
14
13
12
11
10
9
8
7
6
0
0.05
0.1
0.15
0.2
35
30
25
20
15
10
5
0
0.02
0.04
0.06
0.08
0.1
1
0.8
0.6
0.4
0.2
0
0
0.2
0.4
0.6
0.8
1

S6

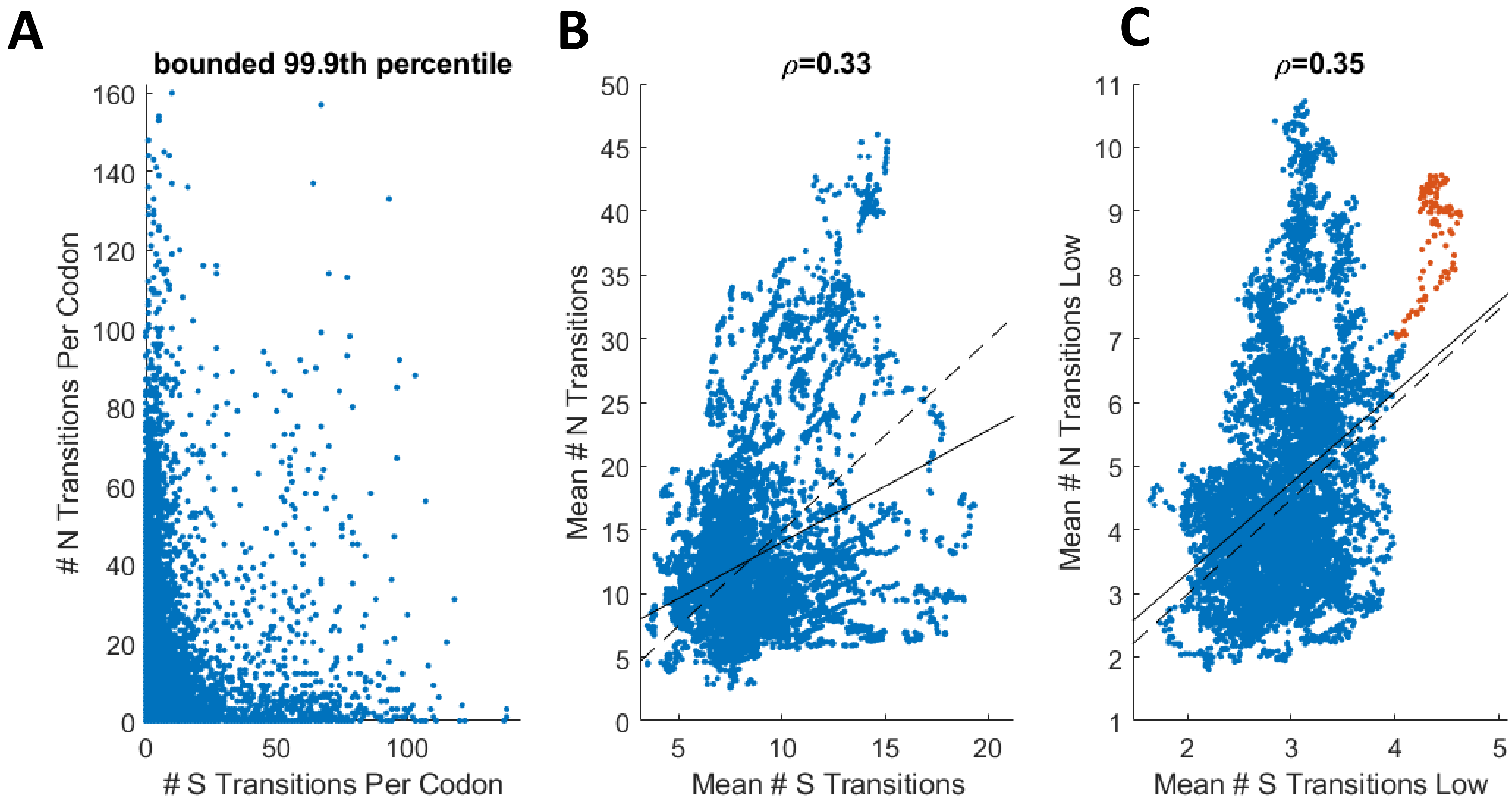
A
bounded 99.9th percentile
### N Transitions Per Codon
### S Transitions Per Codon
0
20
40
60
80
100
120
140
160
0
50
100
B
ρ=0.33
Mean # N Transitions
Mean # S Transitions
0
5
10
15
20
25
30
35
40
45
50
5
10
15
20
C
ρ=0.35
Mean # N Transitions Low
Mean # S Transitions Low
1
2
3
4
5
6
7
8
9
10
11
2
3
4
5

# S7

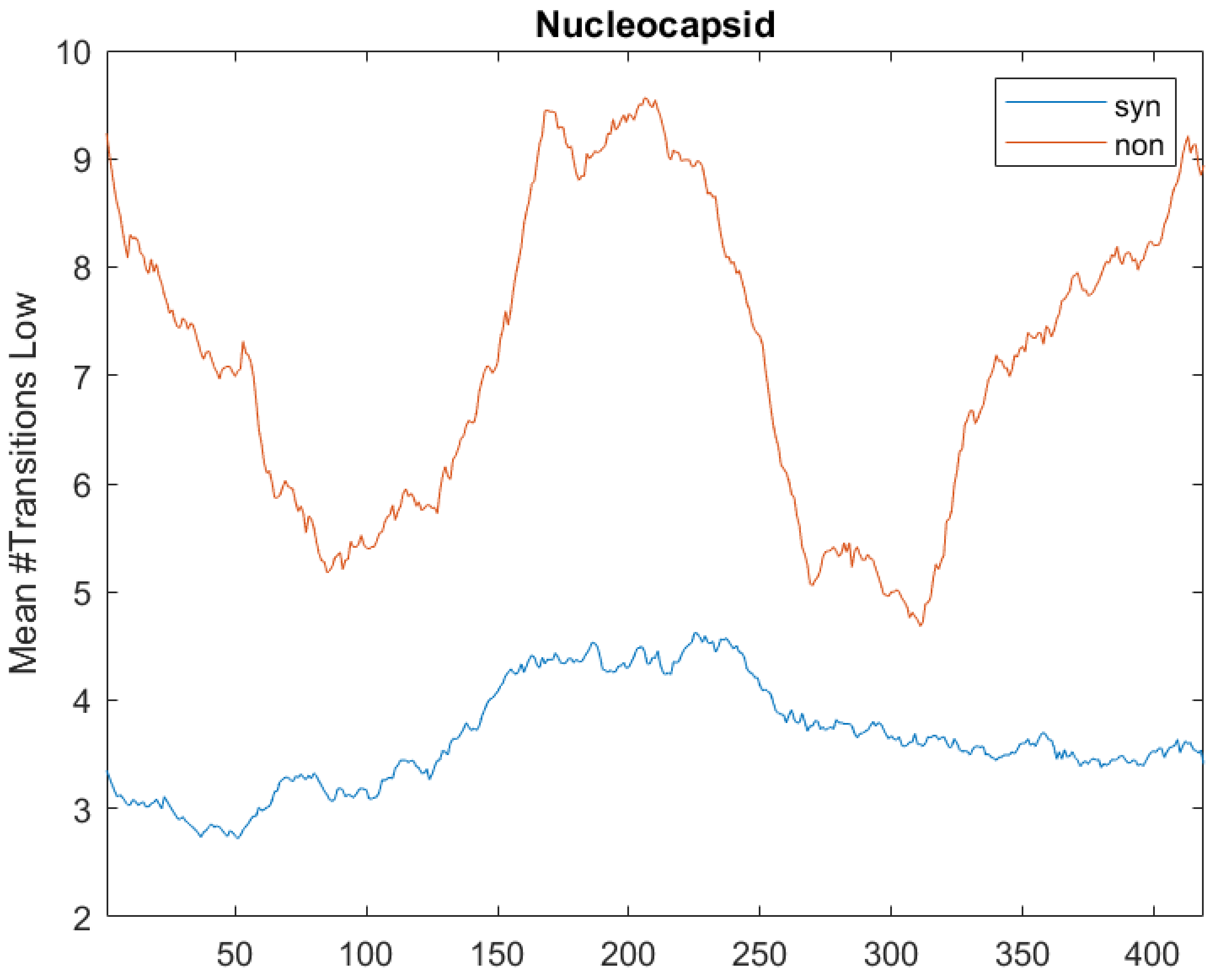
Nucleocapsid
Mean #Transitions Low
10
9
8
7
6
5
4
3
2
50
100
150
200
250
300
350
400
syn
non

**S8**

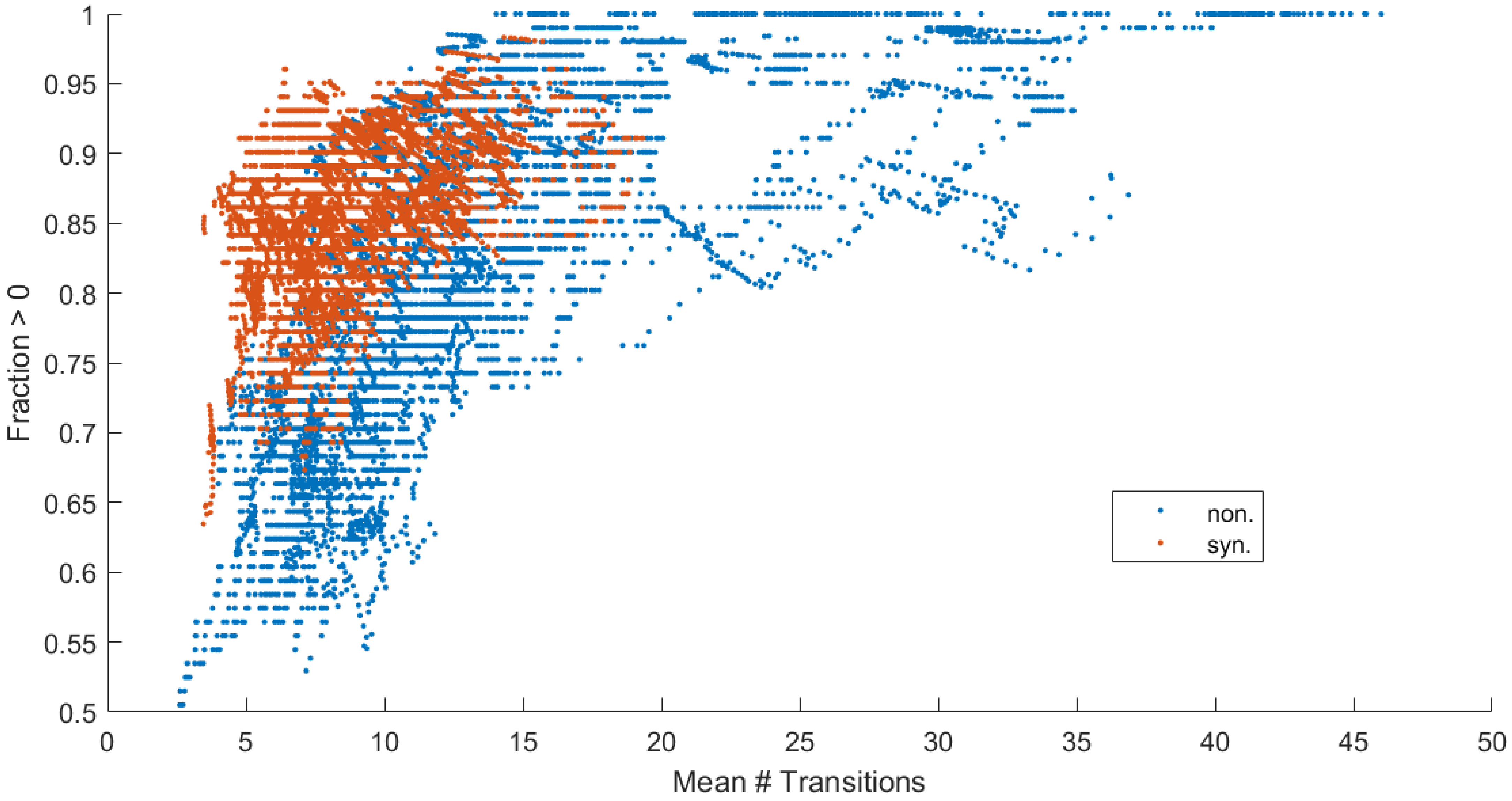

Fraction > 0
1
0.95
0.9
0.85
0.8
0.75
0.7
0.65
0.6
0.55
0.5
0
5
10
15
20
25
30
35
40
45
50
Mean # Transitions
non.
syn.

# S9

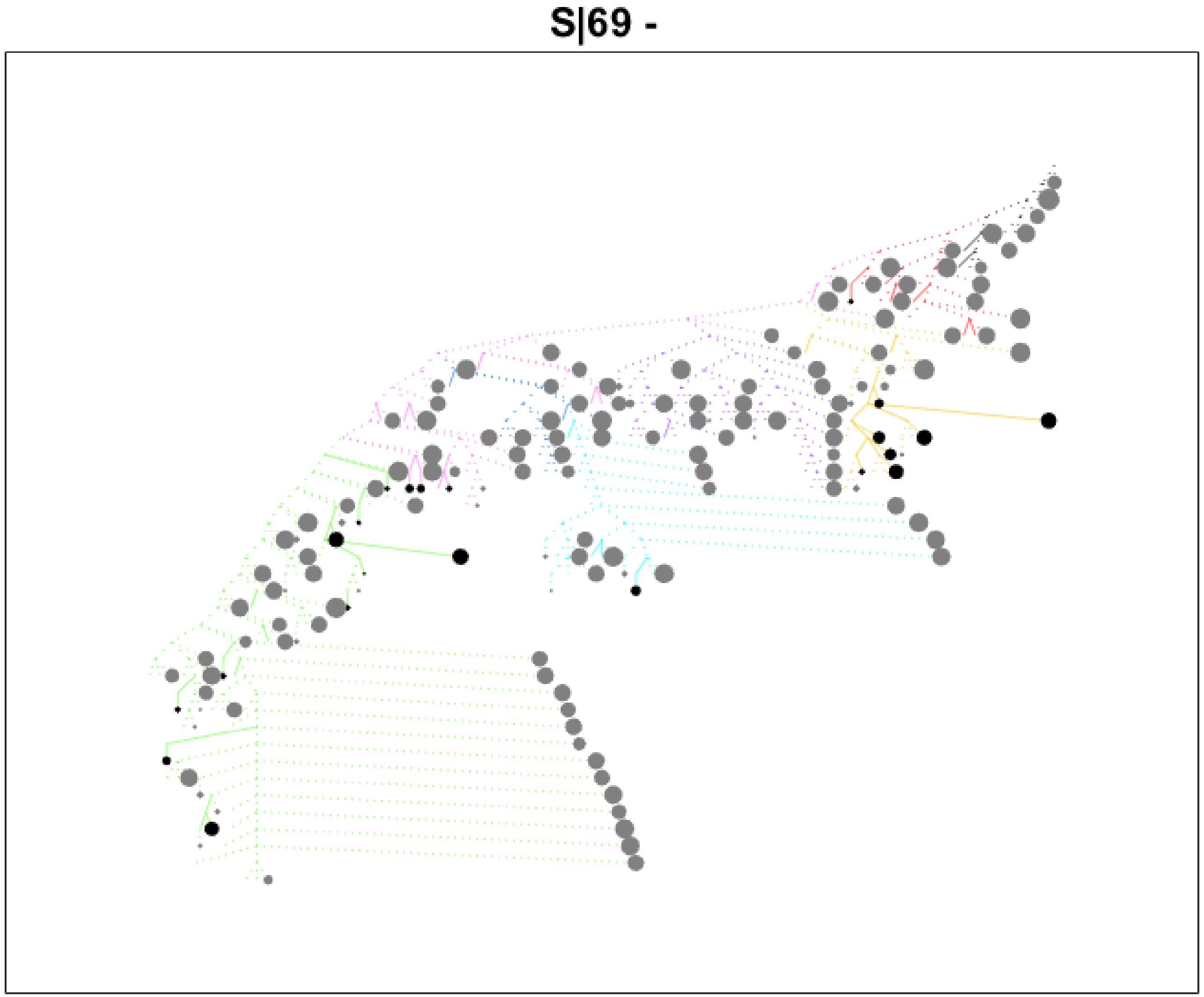

# S10

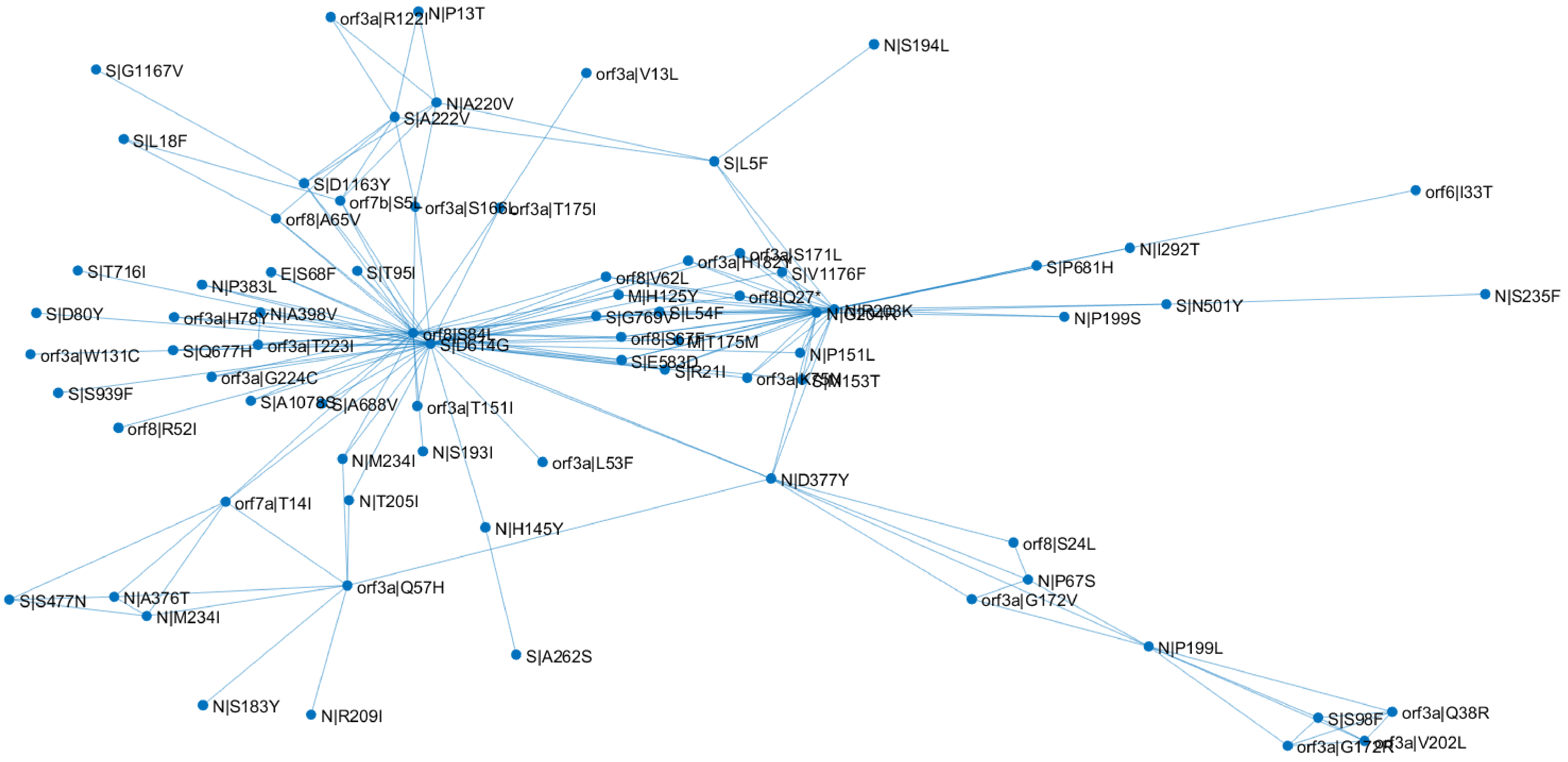

orf3a|R122I
N|P13T
N|S194L
S|G1167V
orf3a|V13L
N|A220V
S|A222V
S|L18F
S|L5F
orf6|I33T
S|D1163Y
orf7b|S5L
orf3a|T175I
orf8|A65V
orf3a|S171L
orf3a|H182Y
N|I292T
S|T716I
E|S68F
S|T95I
orf8|V62L
S|V1176F
S|P681H
N|P383L
M|H125Y
orf8|Q27*
S|D80Y
orf3a|H78Y
N|A398V
S|G769V
S|L54F
N|S235F
S|N501Y
N|P199S
orf8|S84L
S|D614G
orf8|S67F
M|T175M
orf3a|W131C
S|Q677H
orf3a|T223I
N|P151L
S|E583D
S|R21I
orf3a|G224C
S|M153T
S|S939F
S|A1078S
S|A688V
orf3a|T151I
orf8|R52I
N|M234I
N|S193I
orf3a|L53F
N|D377Y
orf7a|T14I
N|T205I
N|H145Y
orf8|S24L
N|P67S
orf3a|G172V
orf3a|Q57H
S|S477N
N|A376T
N|M234I
N|P199L
S|A262S
N|S183Y
N|R209I
S|S98F
orf3a|Q38R
orf3a|G172R
orf3a|V202L

# S11

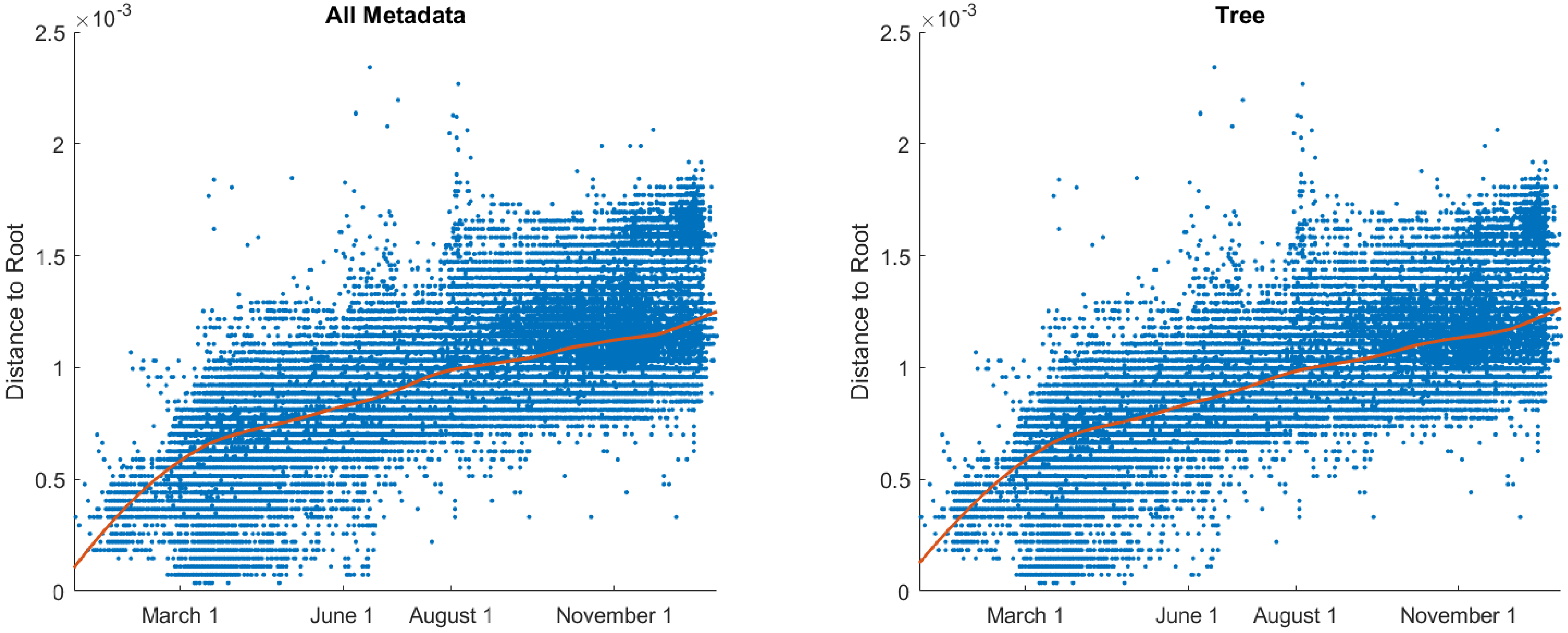
All Metadata
Tree
×10-3
Distance to Root
2.5
2
1.5
1
0.5
0
March 1
June 1
August 1
November 1

**S12**

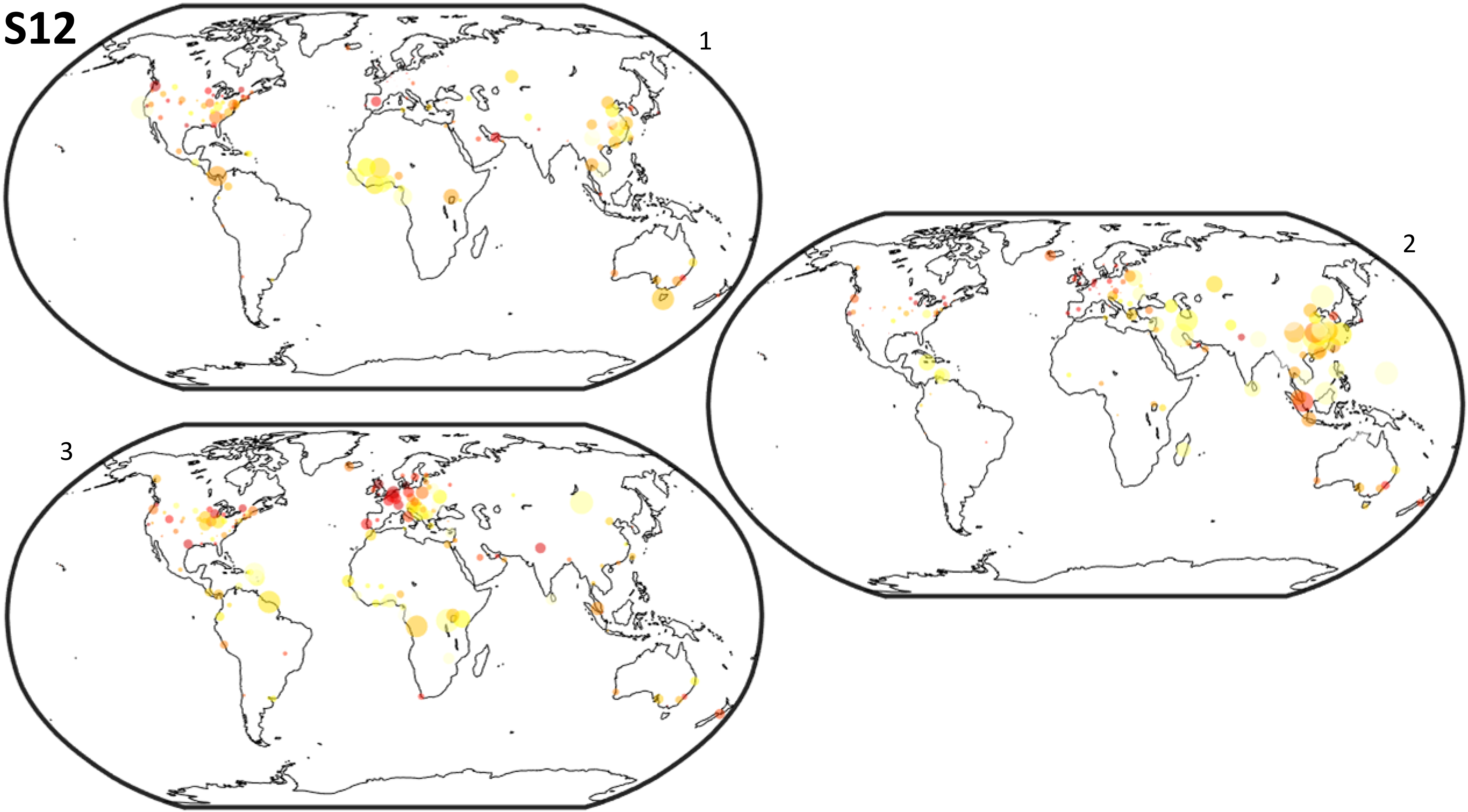

**S13**

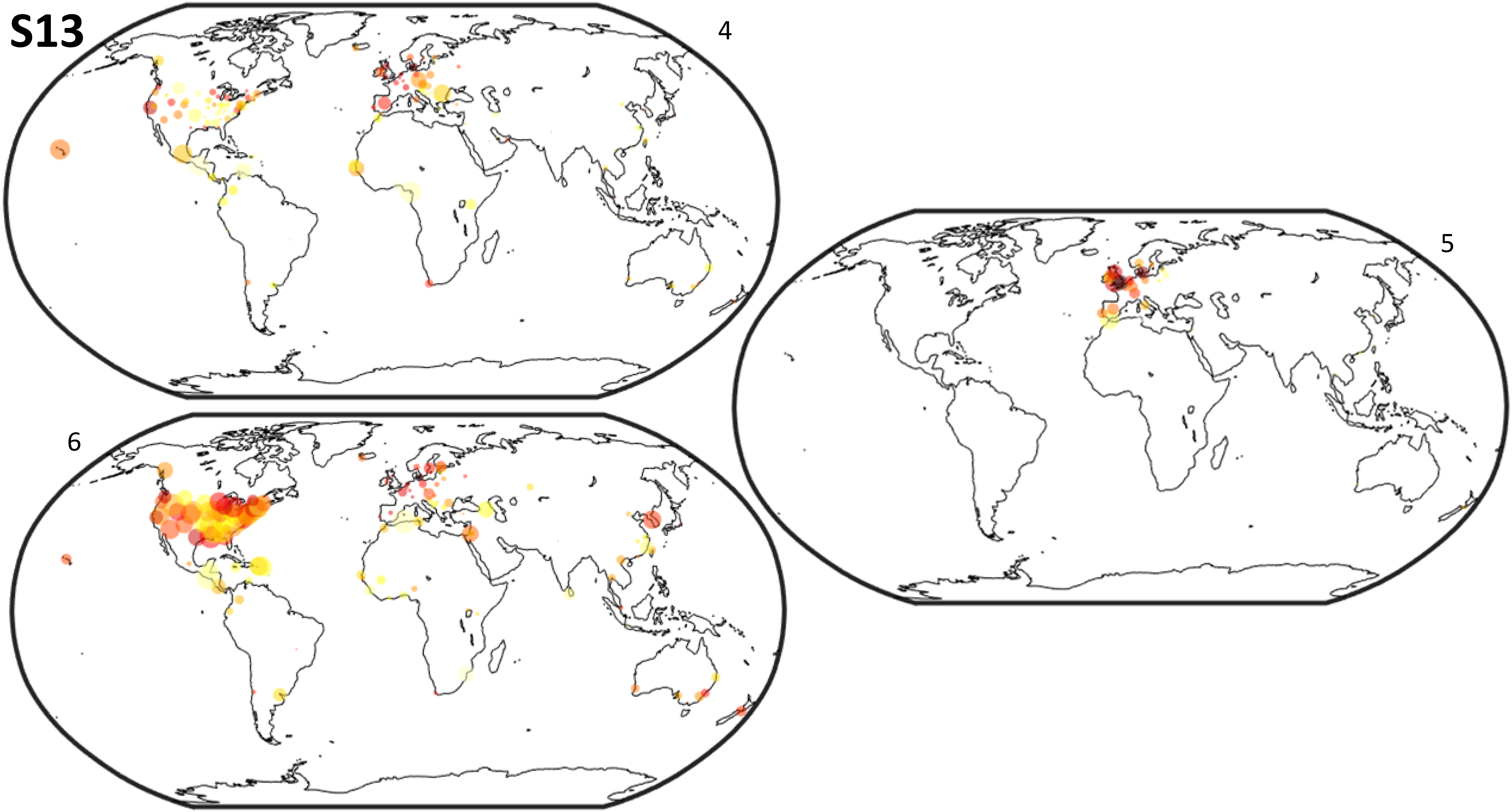

S14

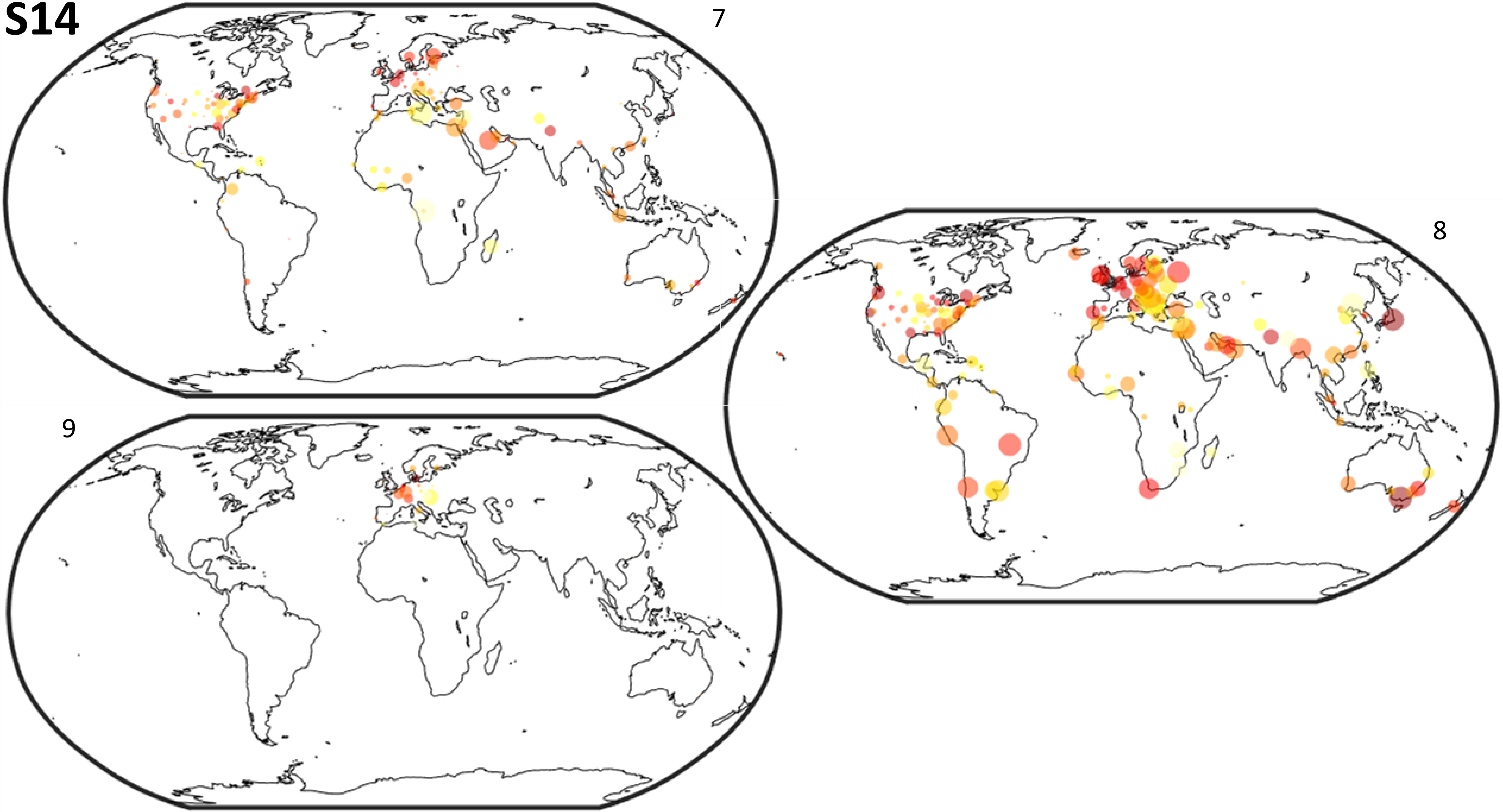

# S15

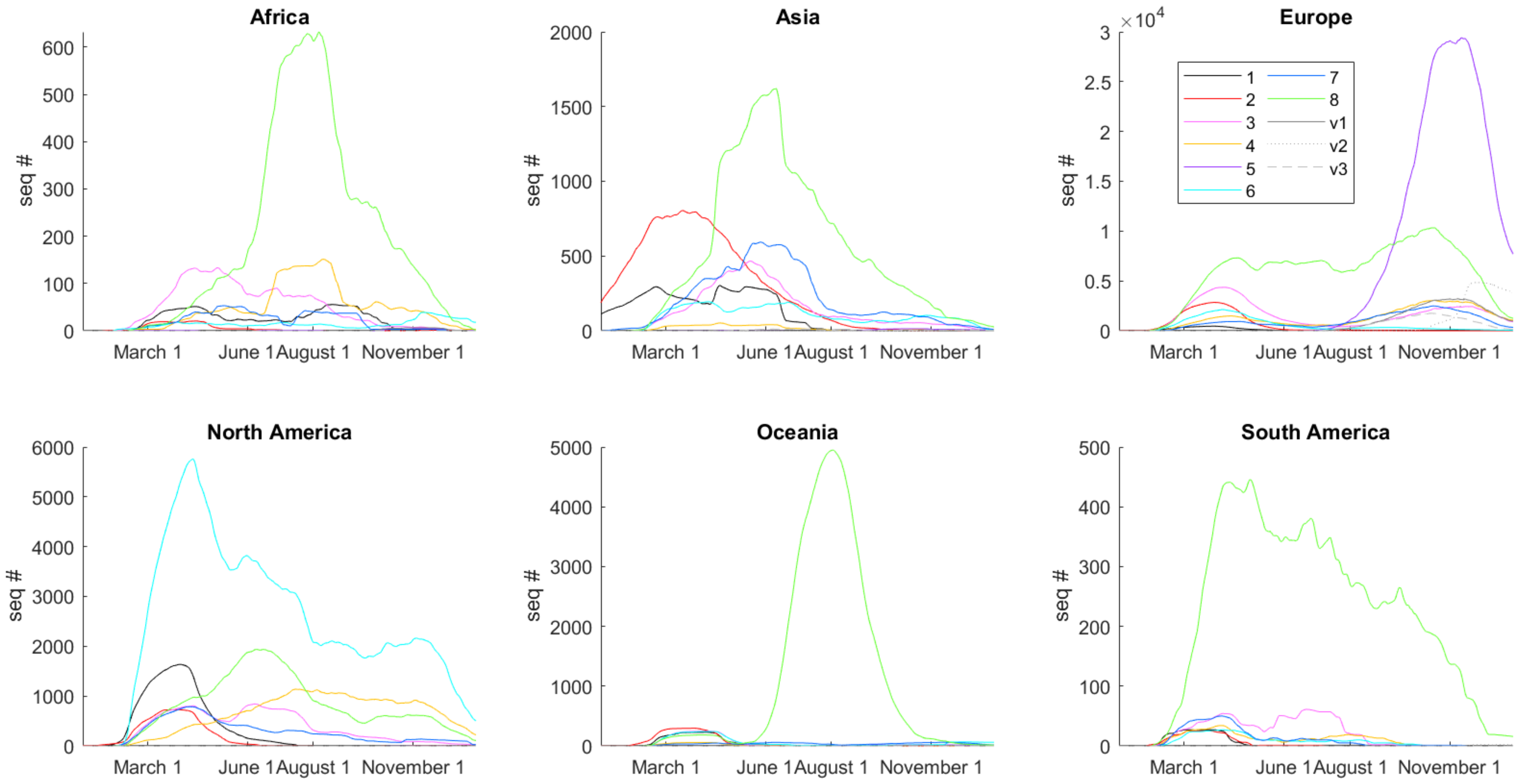
Africa
Asia
Europe
North America
Oceania
South America
seq #
March 1
June 1
August 1
November 1
$\times 10^4$
1
2
3
4
5
6
7
8
v1
v2
v3

# S16

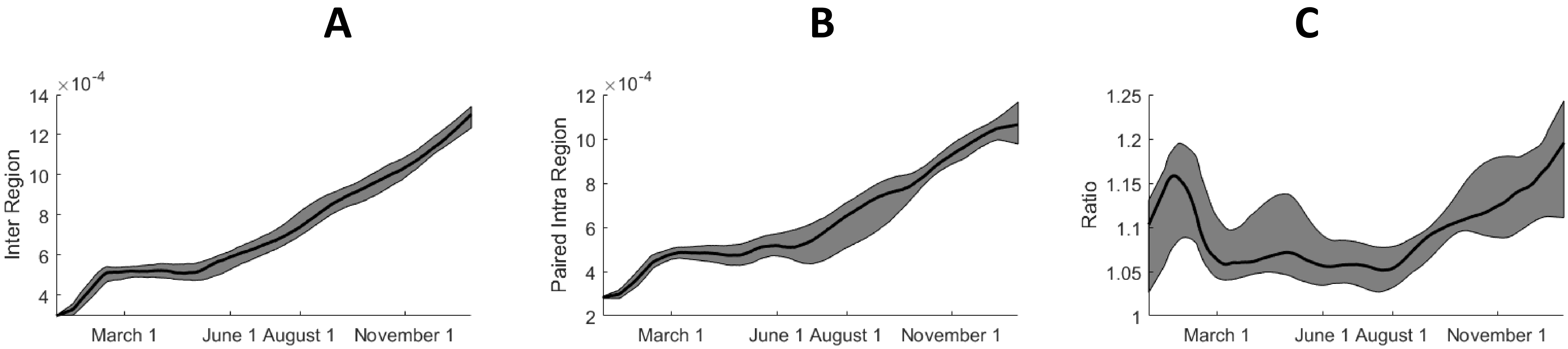
A
B
C
×10^-4
Inter Region
14
12
10
8
6
4
March 1
June 1
August 1
November 1
×10^-4
Paired Intra Region
12
10
8
6
4
2
Ratio
1.25
1.2
1.15
1.1
1.05
1

# S17

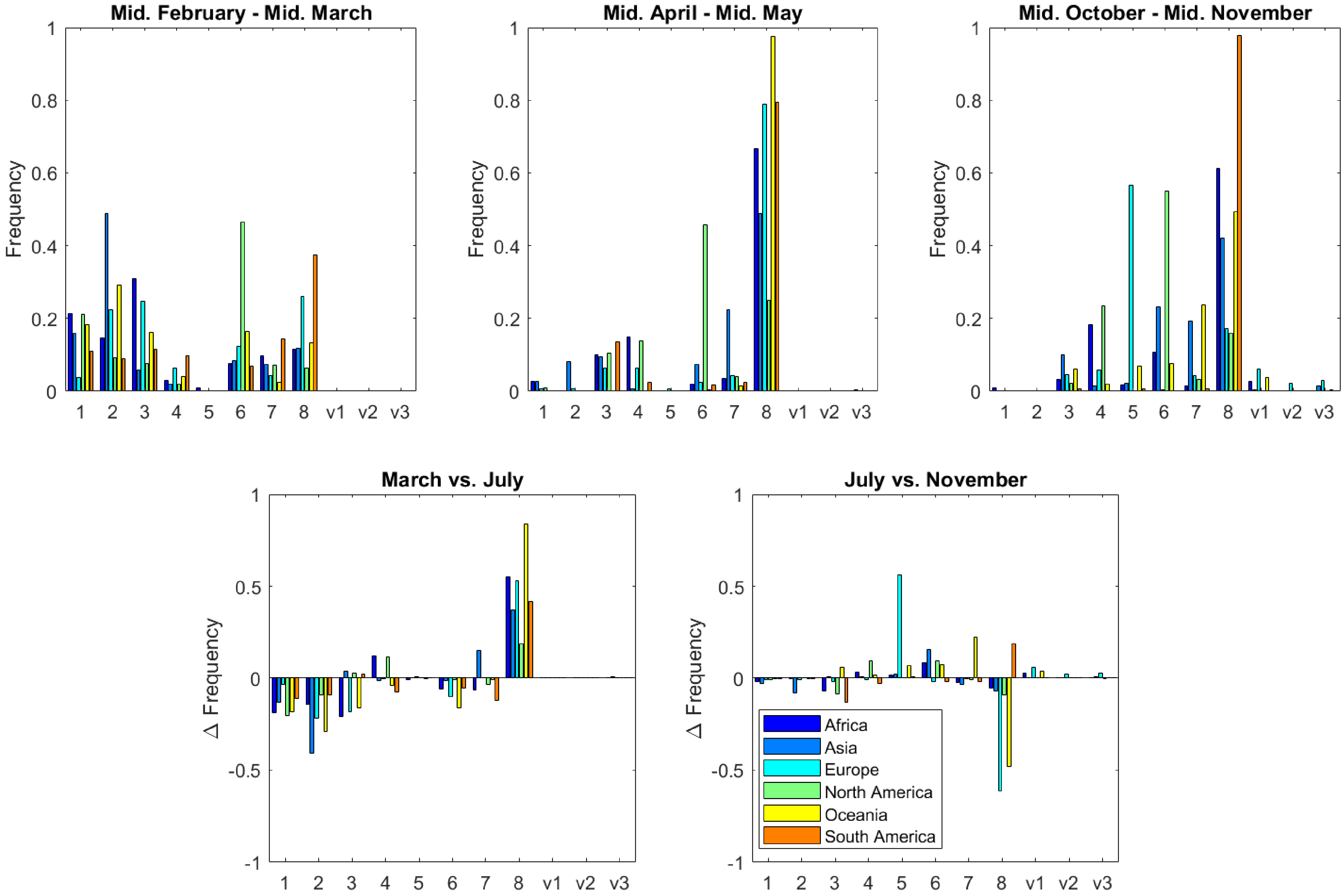
Mid. February - Mid. March
Mid. April - Mid. May
Mid. October - Mid. November
Frequency
0
0.2
0.4
0.6
0.8
1
1
2
3
4
5
6
7
8
v1
v2
v3
March vs. July
July vs. November
Δ Frequency
-1
-0.5
0
0.5
1
Africa
Asia
Europe
North America
Oceania
South America

# S18

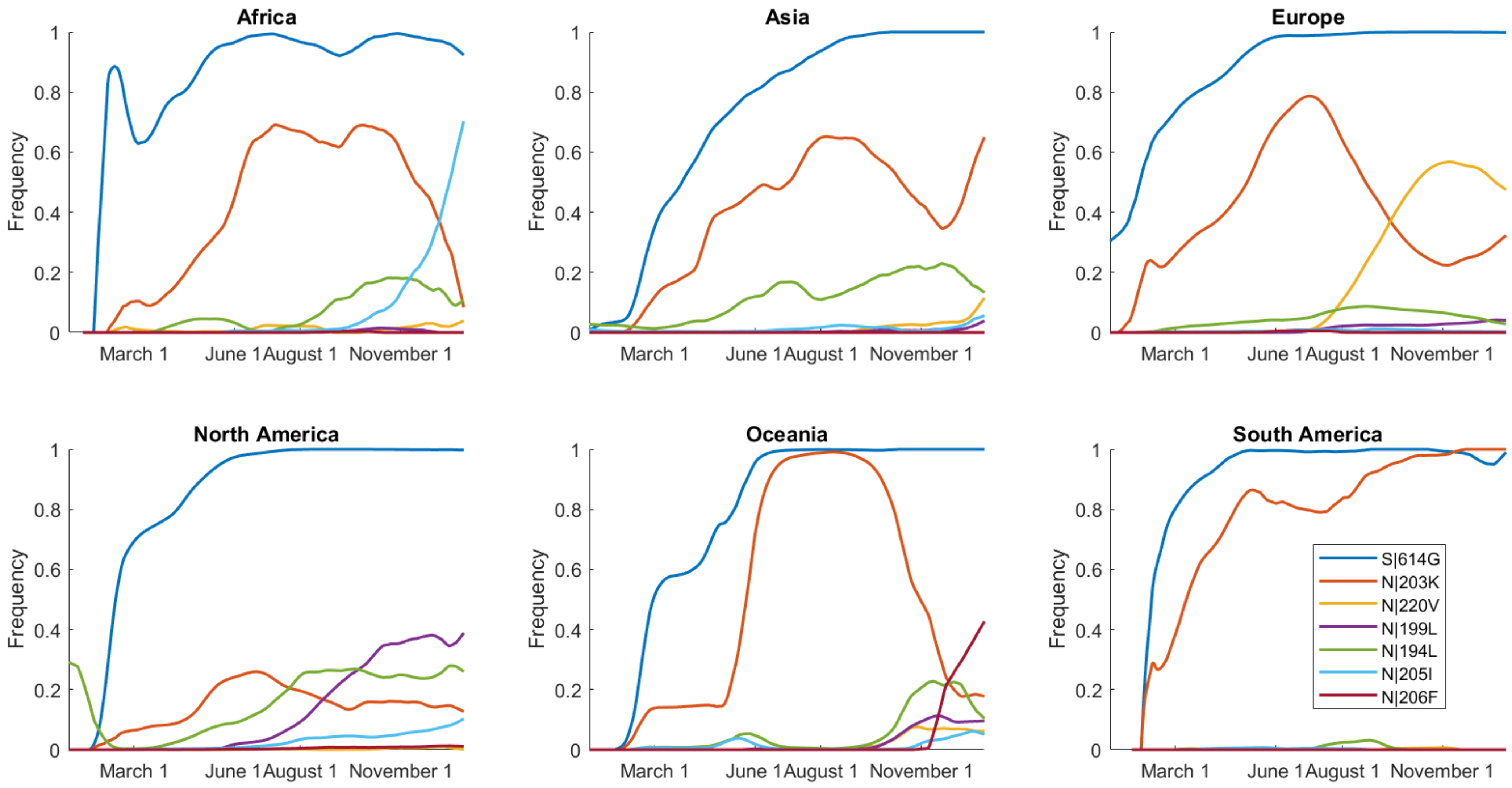
Africa
Asia
Europe
North America
Oceania
South America
Frequency
0
0.2
0.4
0.6
0.8
1
March 1
June 1
August 1
November 1
S|614G
N|203K
N|220V
N|199L
N|194L
N|205I
N|206F

# S19

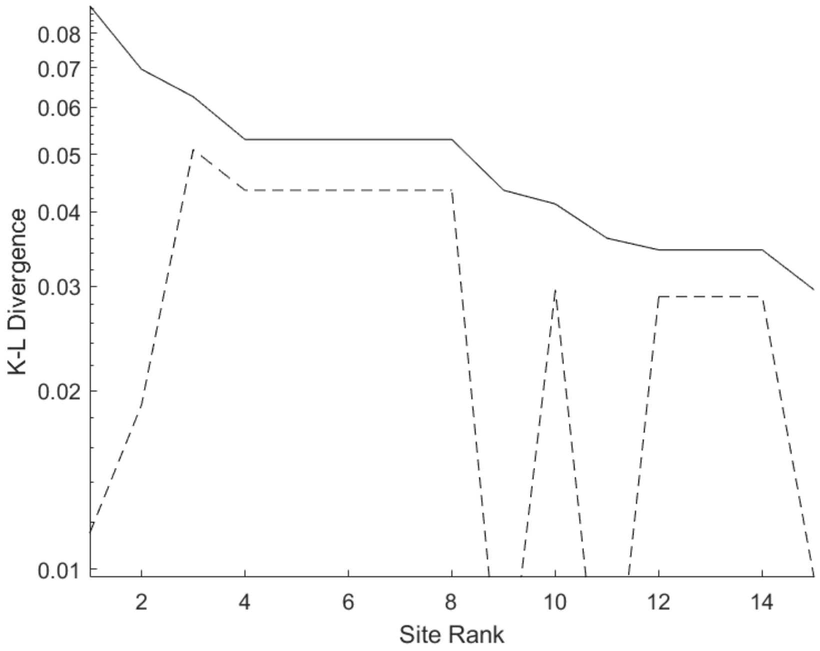


Partition 7, Late 2020, Oceania

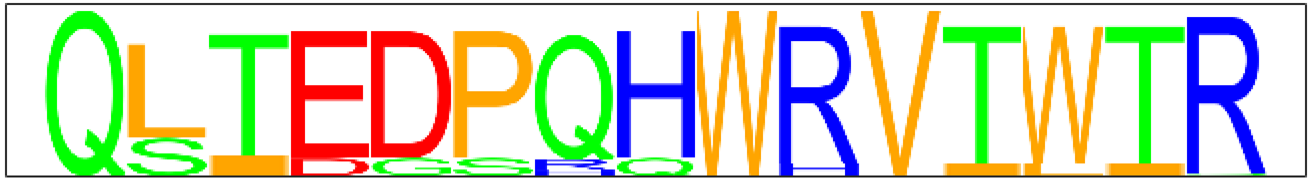

Partition 7, Oceania

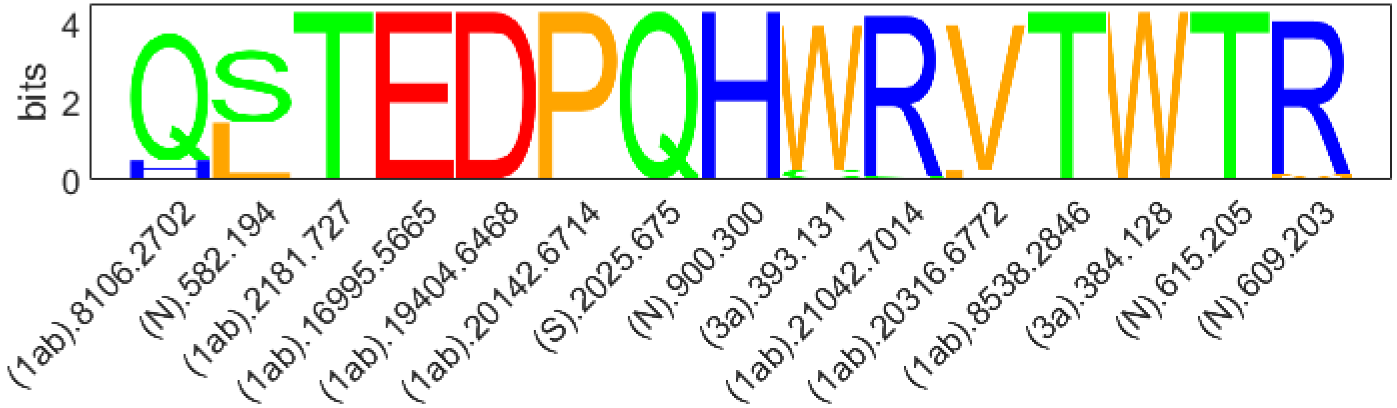

# S20

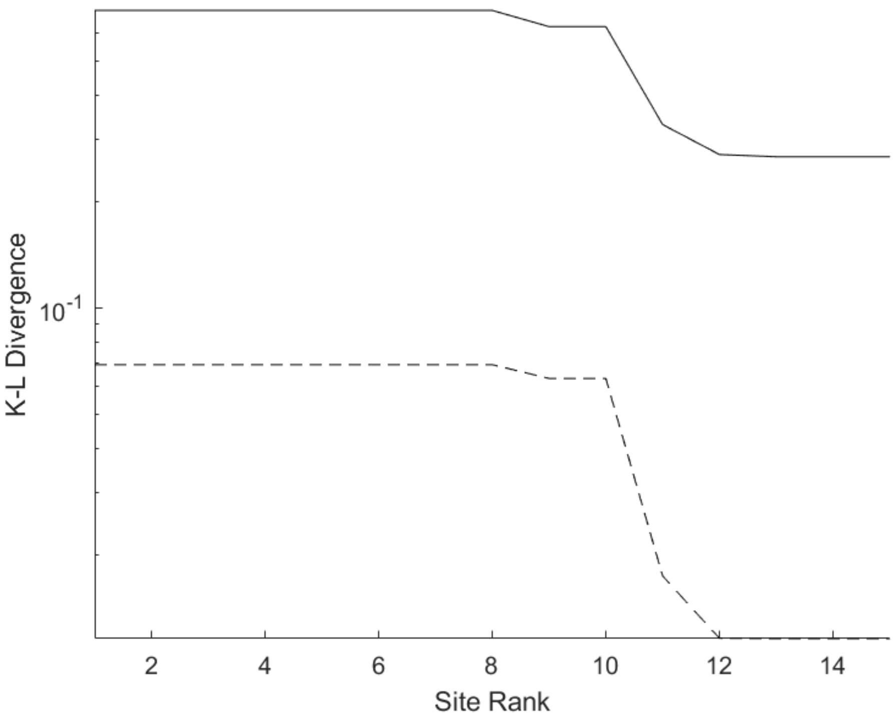


Partition 6, Late 2020, Oceania

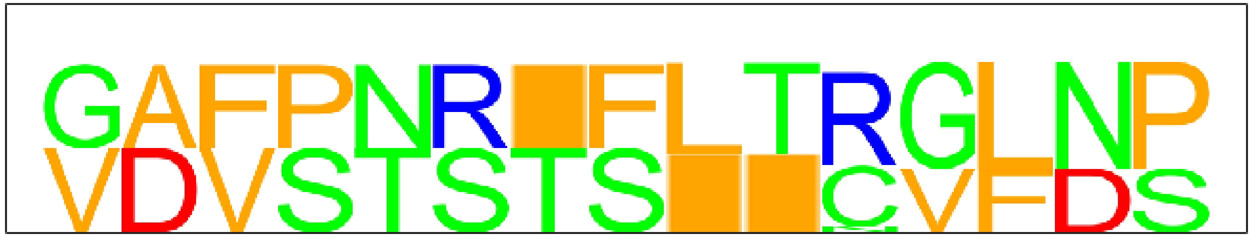

Partition 6, Oceania

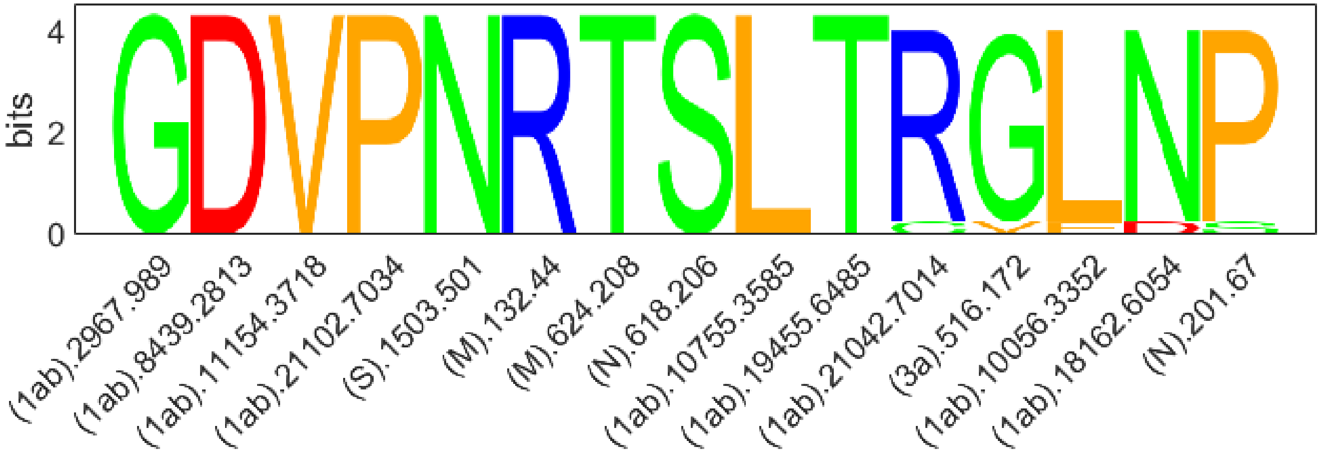

# S21

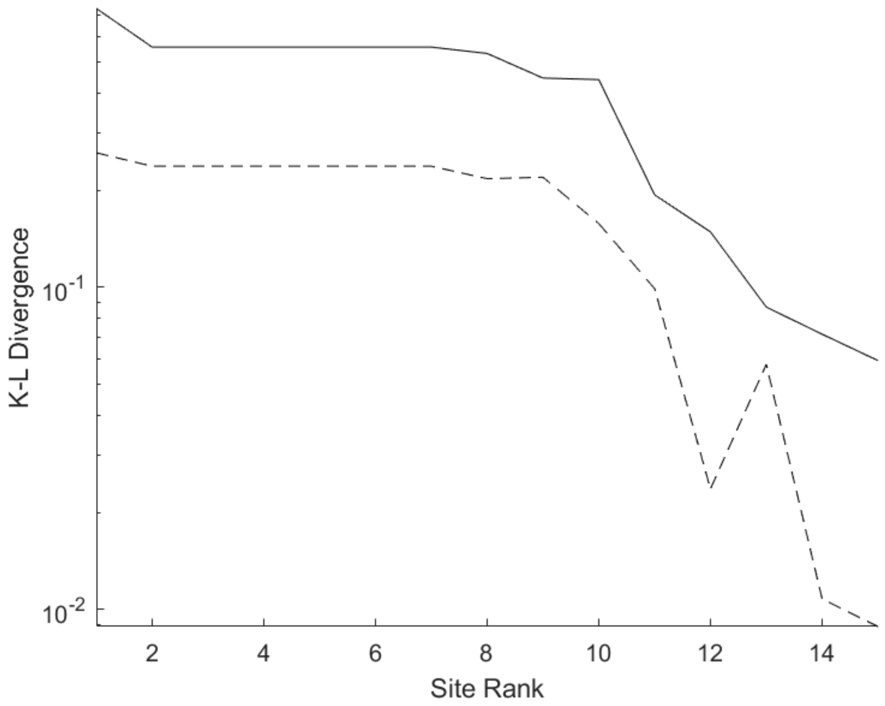


Partition 6, Late 2020, Africa

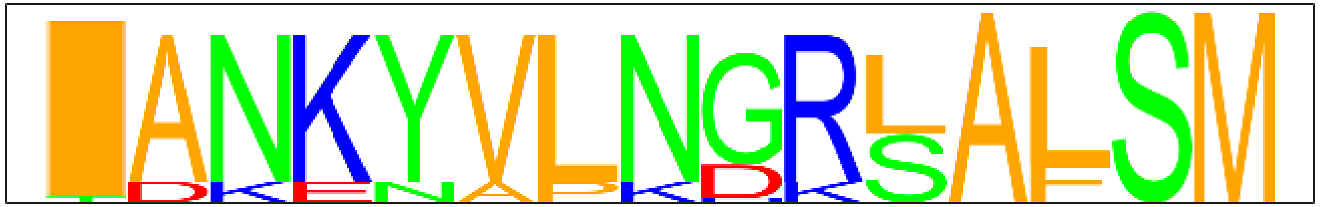

Partition 6, Africa

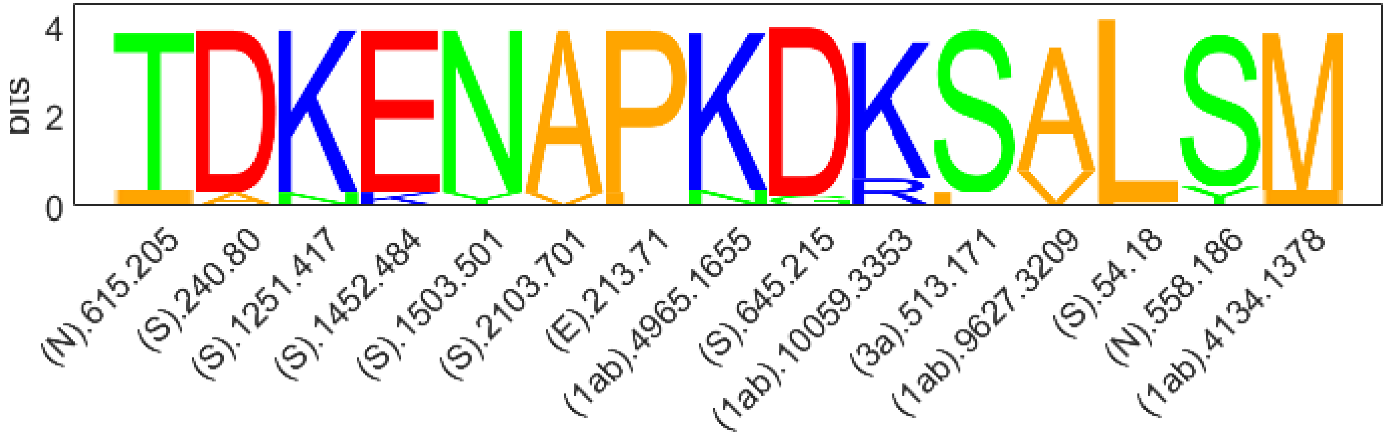

# S22

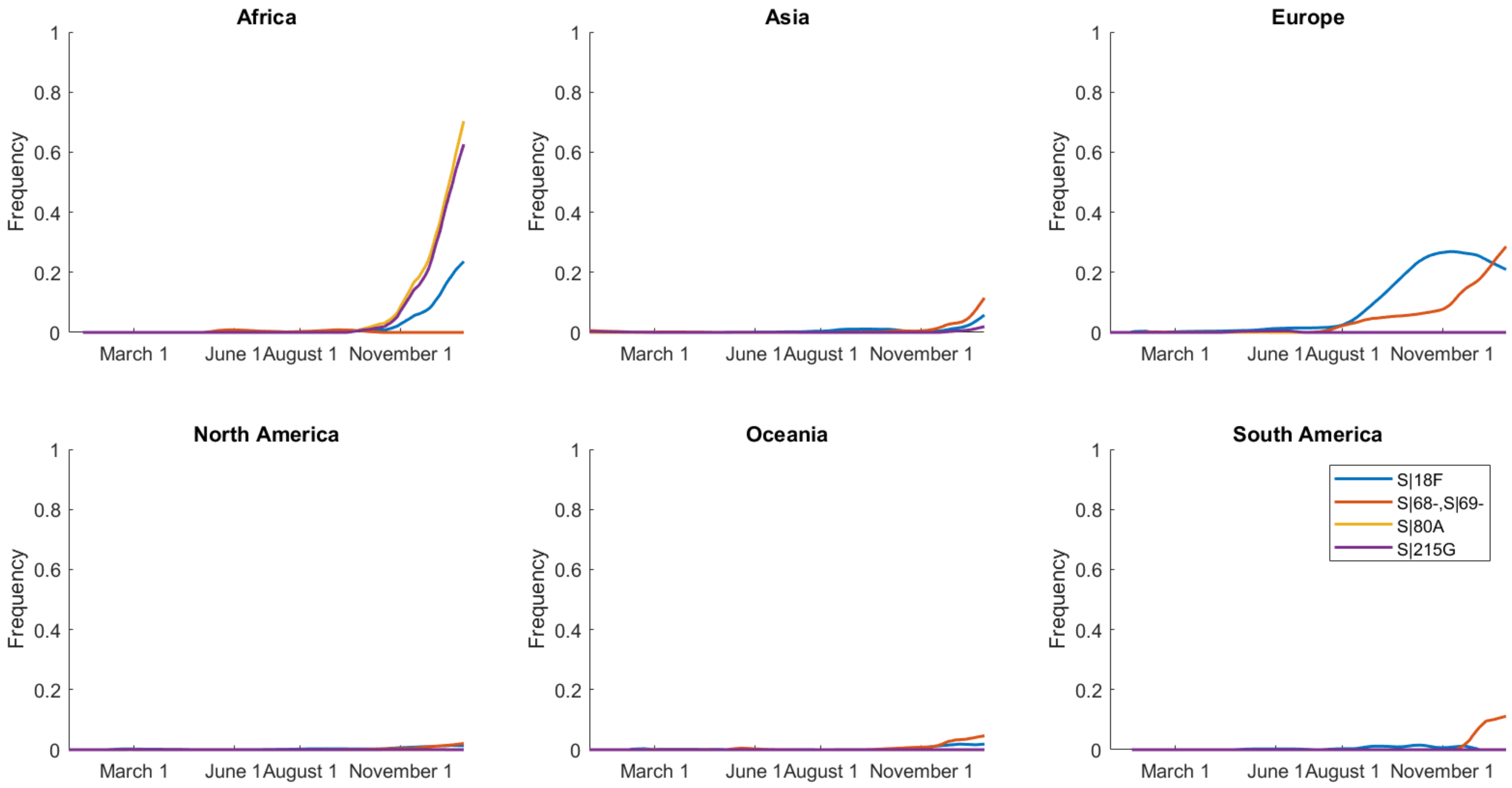
Africa
Asia
Europe
North America
Oceania
South America
Frequency
0
0.2
0.4
0.6
0.8
1
March 1
June 1
August 1
November 1
S|18F
S|68-,S|69-
S|80A
S|215G

# S23

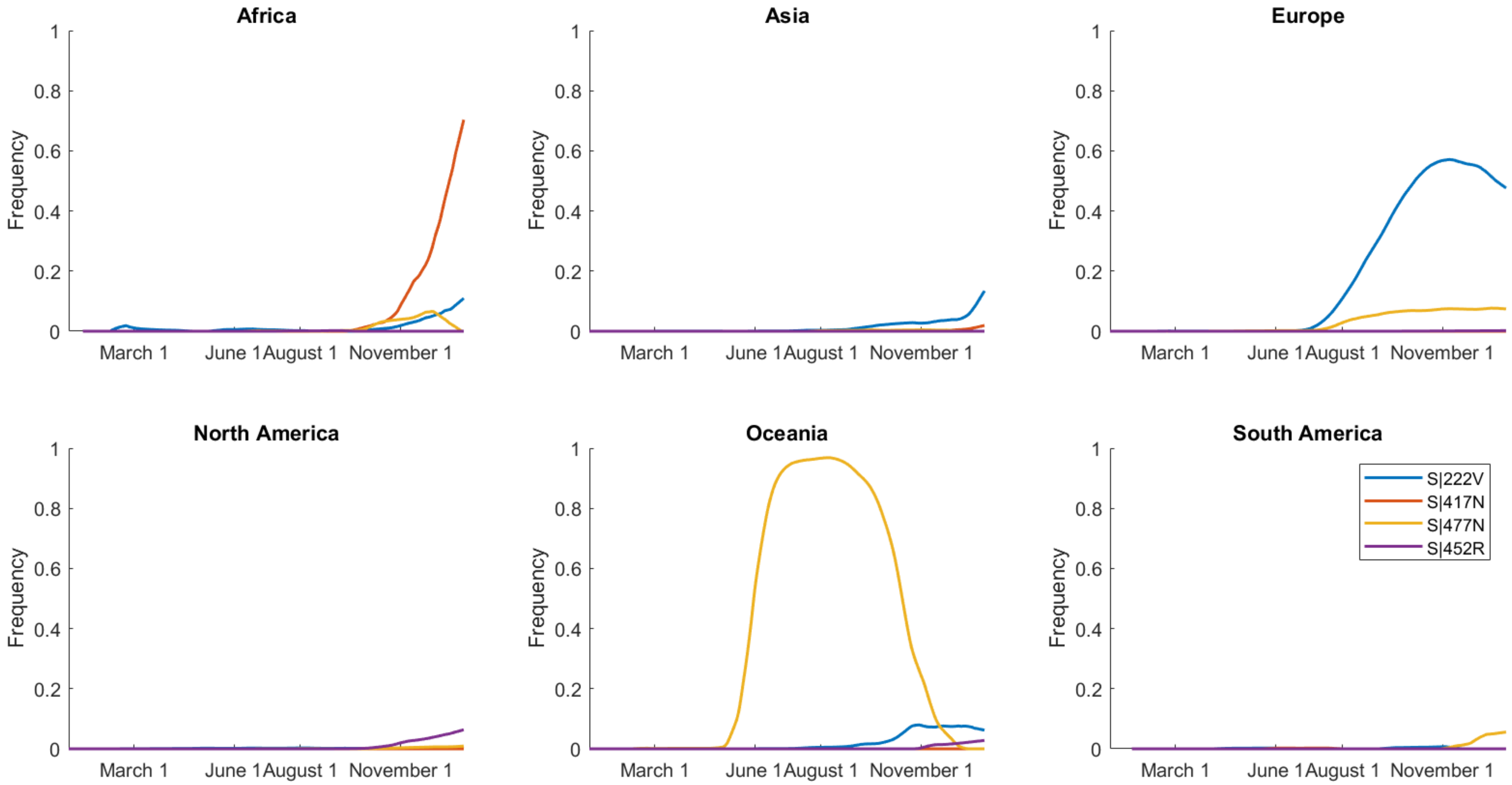
Africa
Asia
Europe
North America
Oceania
South America
Frequency
0
0.2
0.4
0.6
0.8
1
March 1
June 1
August 1
November 1
S|222V
S|417N
S|477N
S|452R

# S24

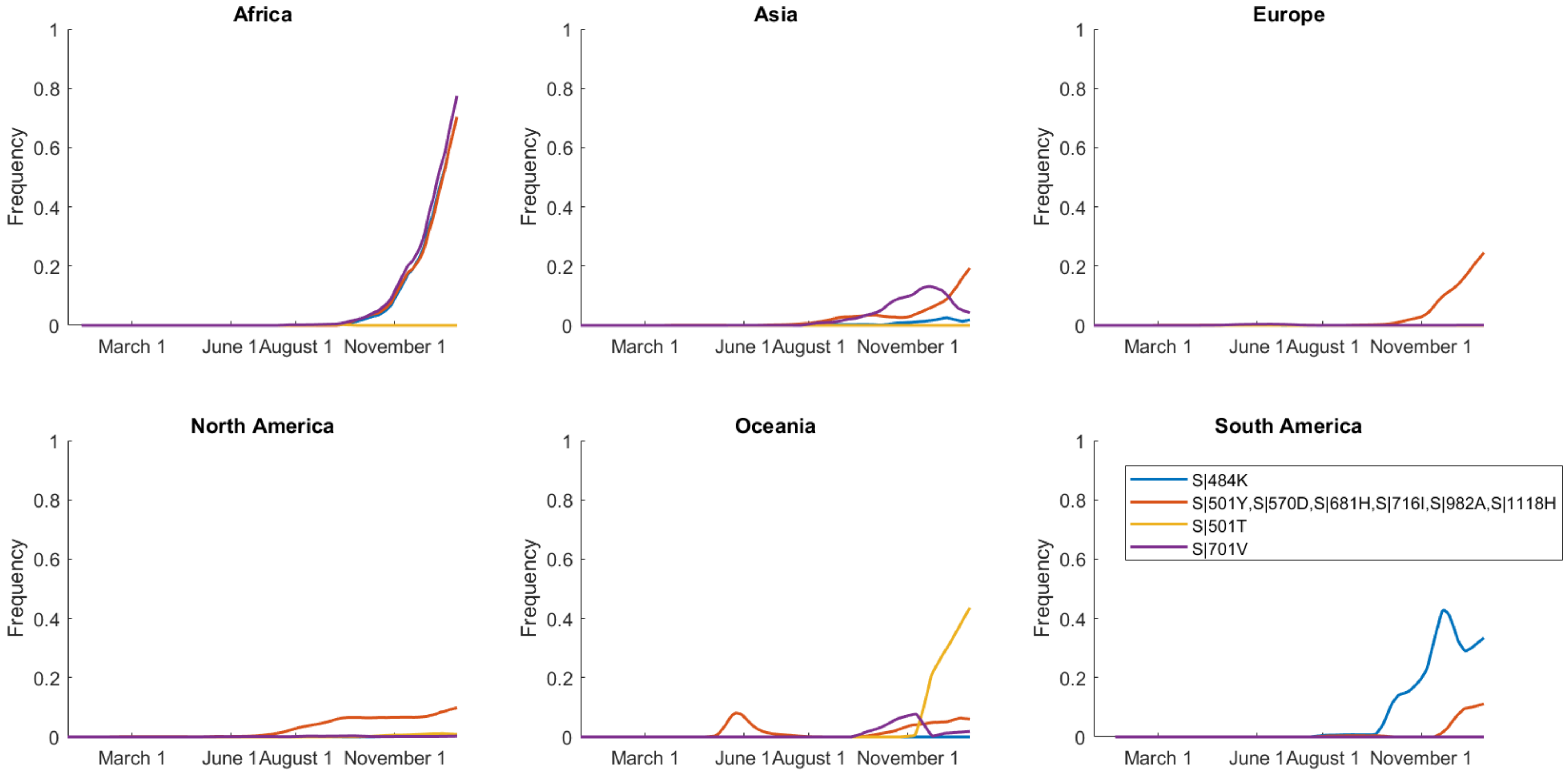

Africa
Asia
Europe
North America
Oceania
South America
Frequency
1
0.8
0.6
0.4
0.2
0
March 1
June 1
August 1
November 1
S|484K
S|501Y,S|570D,S|681H,S|716I,S|982A,S|1118H
S|501T
S|701V

# S25

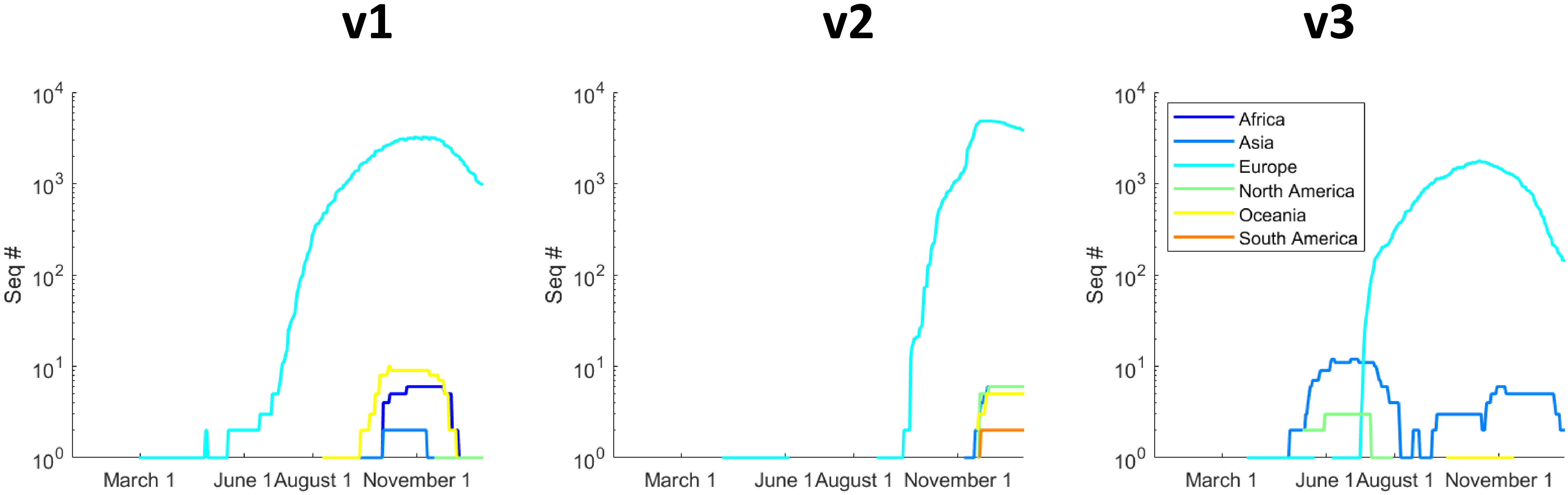

v1
v2
v3
10^4
10^3
10^2
10^1
10^0
Seq #
March 1
June 1
August 1
November 1
Africa
Asia
Europe
North America
Oceania
South America
